# Genome-wide Characterization of Diverse Bacteriophages Enabled by RNA-Binding CRISPRi

**DOI:** 10.1101/2023.09.18.558157

**Authors:** Benjamin A. Adler, Muntathar J. Al-Shimary, Jaymin R. Patel, Emily Armbruster, David Colognori, Emeric J. Charles, Kate V. Miller, Arushi Lahiri, Marena Trinidad, Ron Boger, Jason Nomburg, Sebastien Beurnier, Michael L. Cui, Rodolphe Barrangou, Vivek K. Mutalik, Joseph S. Schoeniger, Joseph A. Pogliano, David F. Savage, Jennifer A. Doudna, Brady F. Cress

## Abstract

Bacteriophages constitute one of the largest sources of unknown gene content in the biosphere. Even for well-studied model phages, robust experimental approaches to identify and study their essential genes remain elusive. We uncover and exploit the conserved vulnerability of the phage transcriptome to facilitate genome-wide protein expression knockdown via programmable RNA-binding protein dRfxCas13d (CRISPRi-ART) across diverse phages and their host. Establishing the first broad-spectrum phage functional genomics platform, we predict over 90 essential genes across four phage genomes, a third of which have no known function. These results highlight hidden infection strategies encoded in the most abundant biological entities on earth and provide a facile platform to study them.

---

Bacteriophages (phages) constitute the most abundant and genetically diverse entities on Earth. Interactions of bacteria with an estimated global total of 10³¹ virions significantly shape both human health and environmental ecosystems (*1*). The scale of ecological interactions between phages and their bacterial hosts drives a genetic arms race that continually alters microbial life at the molecular level (*2*). Diversity arising from rapid evolution over large time scales provides a basis for human health innovations such as phage therapy, and a foundation for biotechnology innovations such as Clustered Regularly Interspaced Short Palindromic Repeats (CRISPR) and CRISPR-associated (Cas) protein systems (*3–5*). However, with great genetic diversity comes great unknowns–little is known about the gene content within the overwhelming majority of phages. Compared to their bacterial counterparts, phage genomes encode a far smaller fraction of genes with known or predicted function, constituting one of the largest sources of genetic dark matter (genes of unknown function) in the biosphere (*6*). Although it is possible to bring some of this dark matter to light using classical genetic techniques, higher-throughput experimental approaches are needed to simplify and expedite characterization of the vastly underexplored genetic content of phage genomes.

Over the last decade, genome-scale CRISPRi has emerged as a primary starting point for probing gene functions in diverse organisms by programmably binding DNA to block transcription with a nuclease-deactivated Cas9 or Cas12 (dCas9/dCas12) and guide RNAs (*7–13*). Systematic extension of this CRISPRi technology to phages has been lacking. Recently, use of dCas12 has been reported for two model dsDNA phages in genome-wide arrayed assay format (*14*). However, establishing CRISPRi technology as a seamless and high-throughput strategy across diverse phages made up of different genetic content, genome organization and lifestyle remains challenging. Phages constitute four Baltimore classes (ssRNA, dsRNA, ssDNA, and dsDNA) (*15*). They implement complex and diverse DNA modifications (*16*, *17*), and some employ advanced genome compartmentalization programs (e.g. two stage injection or the phage nucleus) (*18–20*). Several of these attributes hinder or preclude genomic accessibility to DNA-targeting enzymes like Cas9 or Cas12 (*21–23*), suggesting that a CRISPR technology altogether bypassing DNA targeting might be ideally suited as a phage functional genomics platform.

Given the recently reported vulnerability of phage transcripts during infection (*23*), we posited that the RNA-guided RNA-binding protein dRfxCas13d (HEPN-deactivated *Ruminococcus flavefaciens* Cas13d) (*24*) could be applied as a universal tool for programmable protein expression inhibition. By targeting and binding RNA, dCas13d could circumvent diverse phage genome protection strategies while avoiding the bacteriostatic effects of HEPN-mediated, trans-RNA cleavage encoded by wild-type Cas13 (*25*). Here we present CRISPR interference through antisense RNA targeting (CRISPRi-ART) as a robust method to reduce or eliminate protein expression using dCas13d. We first define CRISPRi-ART design rules, uncovering the susceptibility of ribosome binding sites (RBSs) to dCas13d-targeting in *E. coli*. Then we apply this knowledge through implementation of genome-scale CRISPRi-ART screens in *E. coli* and a diverse set of coliphages. Genome-wide analysis of phage gene fitness efficiently illuminates phage-encoded dark matter critical for infection.

## Targeting dCas13d to bacterial RBSs represses protein expression

Initial experiments showed that targeting dCas13d to the RBS of the *rfp* transcript in *E. coli* reduces RFP production, suggesting that dCas13d can be programmed to compete with the ribosome and impair translation initiation (Fig. 1A) (*24*, *26*). To determine the versatility of this protein repression platform, we tested the programmability and multiplexability of CRISPRi-ART by selectively knocking down the expression of genomically-encoded RFP and/or GFP using CRISPR arrays. Simultaneously targeting both RFP and GFP achieved 10-20 fold reductions with minor variation in reduction efficiency arising from CRISPR RNA (crRNA) spacer positioning (Fig. 1B). To demonstrate that CRISPRi-ART also could knock down expression of an endogenous protein, we targeted the RBS of the nonessential *E. coli* protein DnaK and observed 82.7% (gRNA1) and 97.8% (gRNA5) reduction in DnaK protein abundance compared to a non-targeting crRNA control (fig. S5). Intriguingly, this decrease in protein level was sometimes accompanied by a partial decrease in mRNA abundance for RFP, GFP, and DnaK (fig. S5). This suggests that CRISPRi-ART may act through a combination of translational repression and mRNA destabilization, consistent with previous evidence that reduced translation leads to decreased transcript levels through the loss of mRNA-stabilizing “protective ribosomes” (*27–29*). An attenuated effect on mRNA abundance was observed in the downstream gene *dnaJ* of the same operon for gRNA5, indicating CRISPRi-ART can impart partial polar effects on downstream genes in a crRNA-dependent manner (fig. S5, supplementary text).

**Fig. 1.**
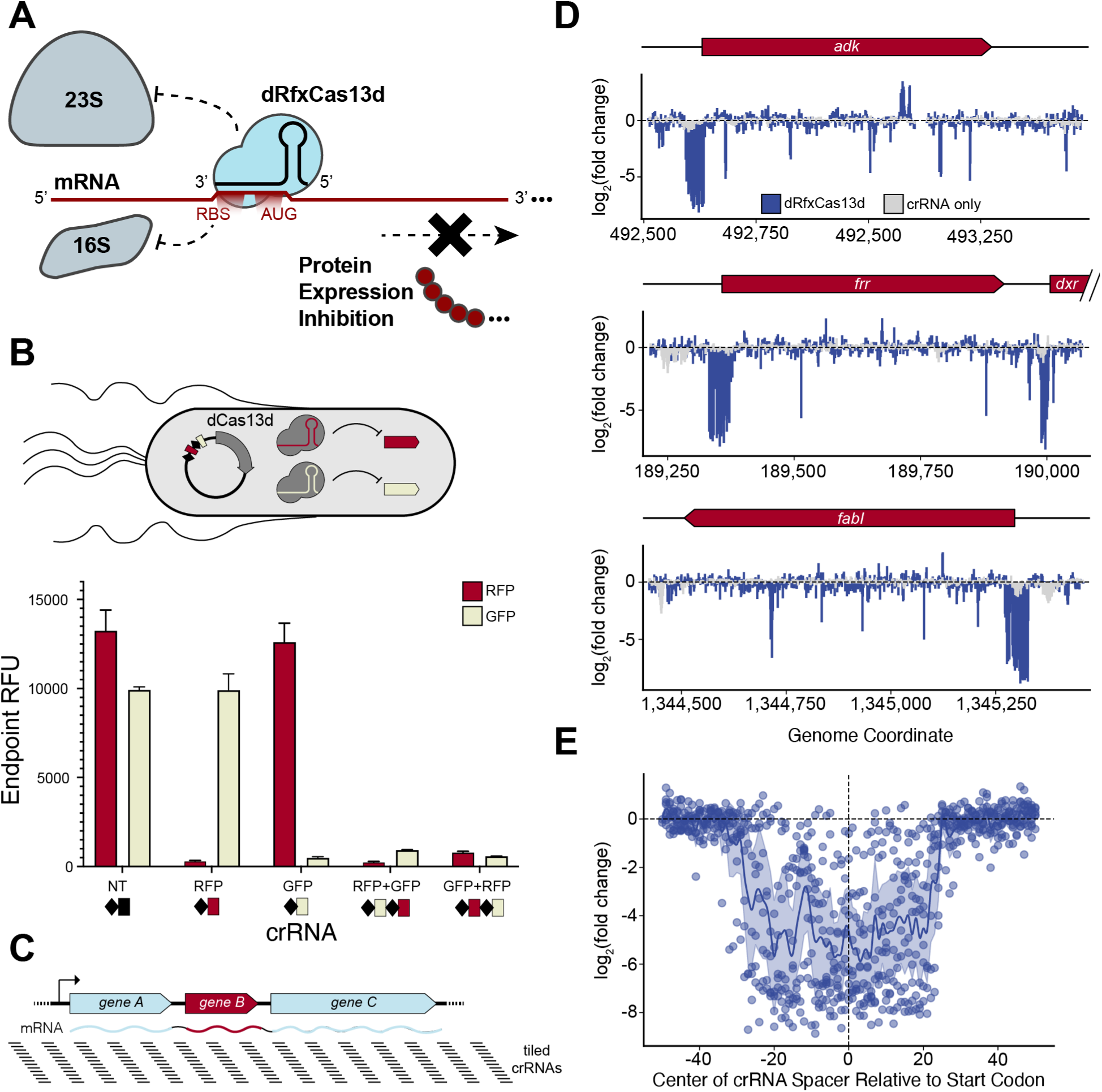
Design rules for dCas13d targeting by exhaustive profiling of *E. coli* essential genes. (**A**)  Overview of CRISPRi-ART. dRfxCas13d binding near the RBS reduces protein expression through inhibition of translation initiation by the 16S ribosomal subunit. (**B**) crRNA arrays were designed against genomically-integrated RFP and GFP. Endpoint relative fluorescence with CRISPRi-ART using a non-targeting (NT) crRNA or a variety of individual and dual crRNAs as indicated. Red and green bars represent RFP and GFP fluorescence, respectively (values are mean +/- SEM with n=12). (**C**) crRNAs were tiled at single nucleotide resolution across transcripts and 100 nt beyond transcript ends, with essential and non-essential gene in red and blue, respectively. (**D**) The measured Log_2_(fold change) of guide abundances targeting various transcripts encoding essential genes is plotted. CRISPRi-ART-dependent fitness effects are presented in blue, and crRNA-only controls in light gray. (**E**) The observed Log_2_(fold change) of crRNA abundances targeting the RBS region of 9 essential genes is compiled into a single plot. Highlighted is the 100 bp surrounding the start codon. Average of values at each nucleotide position is plotted across the region, along with a 95% confidence interval.

Next, we systematically identified regions within mRNA transcripts that are susceptible to CRISPRi-ART. The PAMless nature of dCas13d (*30*) provides a unique opportunity to probe bacterial and phage gene function at single nucleotide resolution, in contrast to PAM-requiring dCas9 and dCas12. Exploiting this advantage, we constructed a pooled, single nucleotide resolution crRNA library tiled across 16 *E. coli* transcripts containing at least one essential gene (Fig. 1C). The 29,473 crRNA library was transformed, induced over 15 doublings, and deep sequenced. This competitive growth assay revealed a major dCas13d-dependent fitness defect for crRNAs binding proximal to the start codon of 9 targeted essential genes (data S6), which we hereby refer to as the RBS (Fig. 1D). Analyzing the fitness measurements (Methods) of all crRNAs for this essential gene set revealed that a tract of 70 nucleotides centered around the RBS was highly susceptible to targeting (Fig. 1E), often producing fitness defects greater than 100-fold. For some of the targeted essential genes, this characteristic RBS-centered fitness defect tract was not observed; we noted that a distinguishing feature of these genes was a markedly lower protein synthesis rate (*31*), suggesting that CRISPRi-ART might be more effective at targeting highly expressed proteins. Thus, by designing crRNAs to target RBSs, we postulated that CRISPRi-ART could efficiently perform genome-wide interrogation using only a few guides per targeted gene.

## CRISPRi-ART enables genome-scale interrogation of the host transcriptome

Identifying the susceptible region surrounding the RBS enabled straightforward compression of genome-scale libraries by targeting crRNAs only to the start of each gene. Specifically, we employed 7 crRNAs coarsely tiled in 5 nt increments across the 35 nt tract proximal to the start codon of each protein-coding sequence in *E. coli* (Fig. 2A, S1, S2). Subsequently, *E. coli* expressing the transformed genome-wide pool of 31,902 antisense crRNAs was cultivated in various media compositions chosen to confer clear conditional fitness measurements for specific genes (Fig 2B) (*32*).

**Fig. 2.**
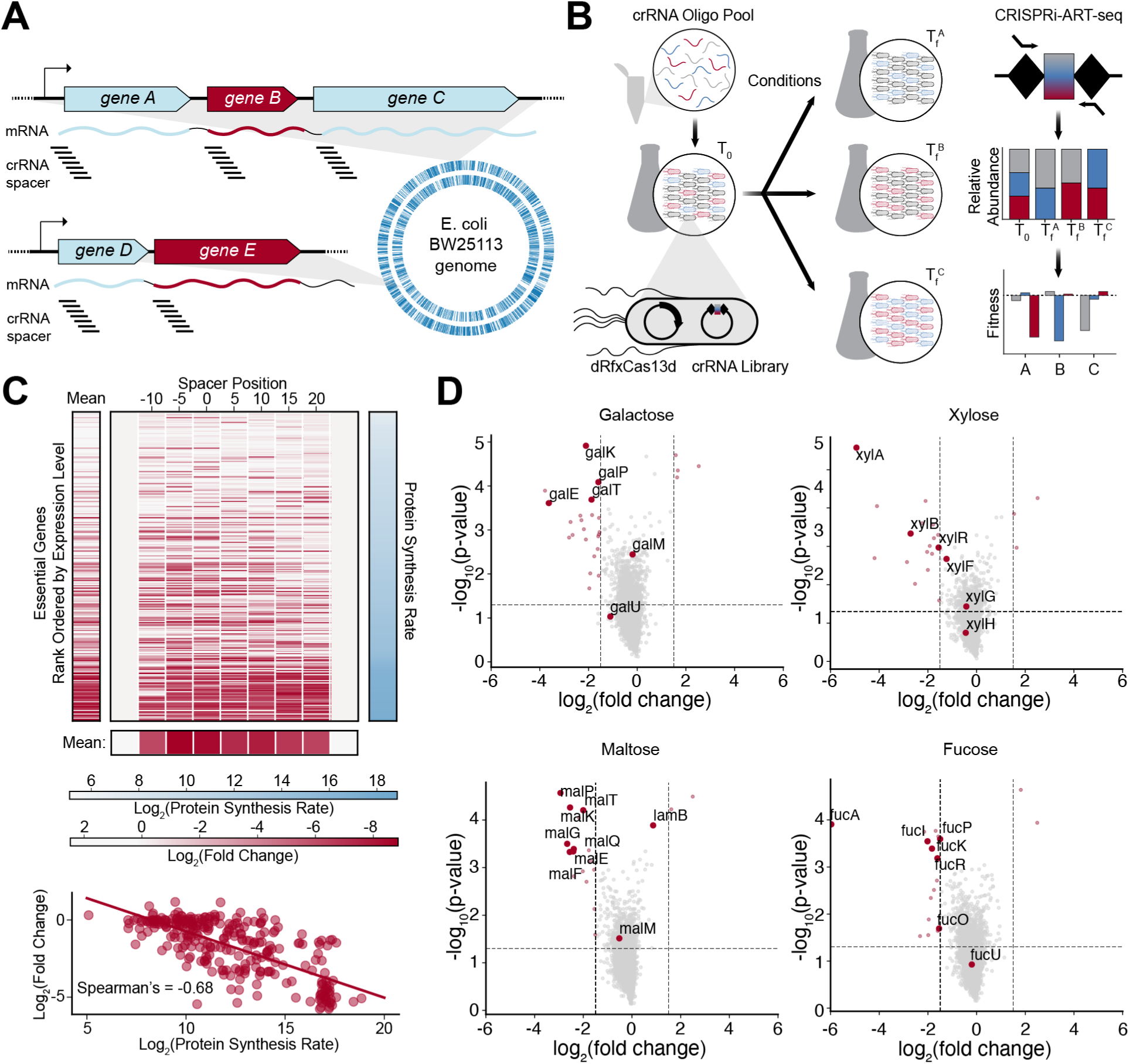
dCas13d interrogates coding elements of the *E. coli* transcriptome. (**A**)  An *E. coli* genome-wide dCas13d crRNA library targets the RBS of each CDS with 7 coarsely tiled crRNAs. (**B**) Simplified schematic of conditional screens and workflow. (**C**) Guide knockdown efficacy correlates with target protein synthesis rate. The Log_2_(fold change) of crRNAs targeting the RBS region of canonical *E. coli* essential genes is plotted (center). For each gene, a mean Log_2_(fold change) is shown (left). Genes are rank-ordered by previously measured protein synthesis rates, the value of which is plotted (right) (*31*). Mean Log_2_(fold change) at each crRNA position is shown (bottom). (D) crRNAs targeting carbon assimilation genes exhibit fitness defects when grown on the corresponding sugar as primary carbon source. Significant differentially-regulated genes are indicated in red (using a Log_2_(fold change) +/-1.5 and 2-tailed T-test p-value < 0.05 as a cutoff). Highlighted in larger markers with labels are genes known to participate in the metabolism and import of respective sugars

First, we analyzed dCas13d-induced genome-wide fitness impacts on *E. coli* growth in rich medium (LB). We defined “significantly impacted” genes as those where at least 3 out of 7 tested crRNAs conferred a log_2_(fold change) below -2 (compared to fitness of NT crRNAs in Fig. S4). We compared these significantly impacted genes with those determined to be “essential” by two established approaches used to generate loss-of-function *E. coli* mutants, 1) saturated transposon-directed insertion site sequencing (*33*) and 2) arrayed single gene deletion mutants (Keio collection) (*34*). This analysis revealed that genome-scale CRISPRi-ART successfully knocked down approximately half of the essential *E. coli* proteome (fig. S3). Similar to the single nucleotide-tiling assay, we observed a strong correlation between fitness measurement and synthesis rate (*31*) for essential *E. coli* genes (Fig. 2C, S3), suggesting that highly expressed proteins elicit stronger phenotypes from CRISPRi-ART.

A major advantage of genome-scale loss-of-function libraries is ease of screening a variety of conditions to identify conditionally important genes and ascribe putative functional annotations. With this in mind, we next grew *E. coli* expressing the genome-wide crRNA libraries in minimal medium with a variety of sugars as primary carbon sources (*32*, *35*). As seen in Fig. 2D, crRNAs targeting genes involved in assimilation and catabolism of the respective sugars exhibited significant fitness defects relative to growth on glycerol as a sole carbon source. A gene exhibiting a significant fitness defect is defined as having at least 3 crRNAs exhibiting a Log_2_(fold change) in abundance of < -2. Having established CRISPRi-ART as a genome-scale functional genomics platform, we next turned to bacteriophages, whose strong transcriptional and translational programs would make them well-suited as targets for CRISPRi-ART.

## CRISPRi-ART enables targeted disruption of phage infection

Phages with DNA genomes employ diverse strategies to avoid DNA-targeting bacterial defense systems, but their RNA transcripts are vulnerable during infection (Fig. 3A) (*23*). To test if CRISPRi-ART is capable of probing phage gene essentiality, we challenged CRISPRi-ART-containing *E. coli* with model *E. coli* phage T4. We expected to see no loss of infectivity while targeting phage-nonessential genes and reduction of infectivity following targeting of phage-essential genes (Fig. 3B). We designed CRISPRi-ART crRNAs targeting T4-essential genes (*36*) involved in replication (*gp45*), recombination/repair (*gp32*), DNA modification (*gp42*), and virion structure (*mcp*) as well as a known non-essential gene, *soc*. When phage-essential genes were targeted by CRISPRi-ART, we consistently found inhibition of phage infection with efficiency of plaquing (EOP) reduced ∼100-10000X (Fig. 3C, fig. S6). In contrast, targeting of *soc* yielded no phage inhibition (Fig. 3C, fig. S6), suggesting that reduction in EOP was conferred due to the essentiality of the targeted gene.

**Fig. 3.**
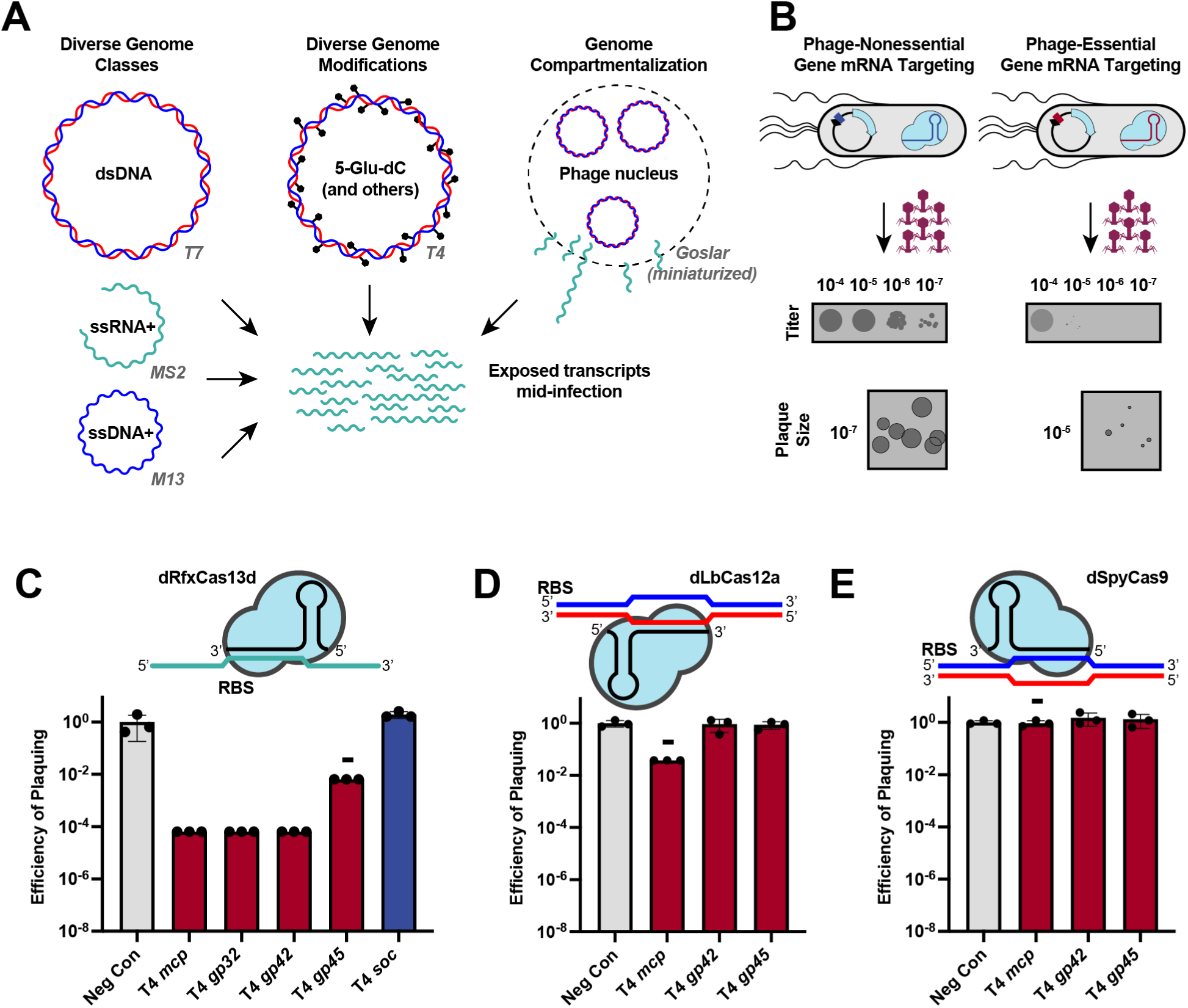
Translational repression provides a simple means to probe phage gene essentiality. (**A**)  Phage-encoded genome protection strategies. Phage genomes can be constituted by ssRNA+ (green), ssDNA+ (blue), or dsDNA (blue/red) molecules (left), heavily modified (center), or compartmentalized (right) with example phages tested in this study. In all cases, phage mRNA (green) is accessible to Cas13-targeting. Phage genomes not drawn to scale. (**B**) Overview of CRISPRi-ART-mediated phage defense. Phage infection is inhibited if dCas13d targets an essential phage protein’s RBS, while infection is productive if the encoded protein is dispensable. Plaque images shown are cartoon illustrations representative of collected data across Figs. 3 and 4. (**C**) EOP assays for CRISPRi-ART-mediated phage defense when targeting phage T4 genes. (**D**) EOP assays for DNA-targeting dCas12a targeting phage T4 genes. (**E**) EOP assays for DNA-targeting dCas9 targeting phage T4 genes. Gray bars represent a negative control crRNA, dark red bars, a known T4-essential gene targeting crRNA, and dark blue bars, a known T4-nonessential gene targeting crRNA. All EOP values represent the average of three biological replicates at 20 nM aTc dCas13d or dSpyCas9 induction or 5nM aTc LbCas12a induction. EOP data are presented as mean ± SD. Minus symbols denote a consistent, ≥4-fold plaque size reduction phenotype if plaques were observed.

To benchmark performance of CRISPRi-ART against previously-established dsDNA-targeting CRISPRi tools for phage inhibition, we repeated phage challenges using dLbCas12a and dSpyCas9 (*7*, *37*). While dSpyCas9 is known to inconsistently bind to modified T4 genomic DNA, dLbCas12a can bind to T4’s modified genome (*21*, *38*). We observed targeting T4-essential genes with dsDNA-targeting dLbCas12a (Fig. 3D, S7) or dSpyCas9 (Fig. 3E, S8) resulted in minimal anti-phage activity. These results, along with a recent successful use of dCas12a to target every gene in Lambda and P1 phages (*14*), suggest that dsDNA targeting CRISPRi platforms are not always reliable, especially for lytic phages with genome protection strategies.

## CRISPRi-ART is broadly effective across E. coli phage phylogeny

To test if CRISPRi-ART is applicable across diverse bacteriophages, we employed dCas13d with 12 diverse coliphages (Fig. 4A). These phages included 3 Baltimore classes (ssRNA+ (MS2), ssDNA+ (M13), and dsDNA); chemically modified genomes (T4, MM02); temperate (Lambda), chronic (M13), and lytic lifestyles; and genome compartmentalization strategies (T5, Goslar) (*15*, *18–20*, *22*, *36*, *39*, *40*). For each phage, we designed two crRNAs (gRNA1 and gRNA4) (fig. S1) targeting an essential gene encoding the major capsid protein (MCP) and measured infection productivity via efficiency of plaquing (EOP) and plaque size (Fig. 4B). For every phage, at least one crRNA caused a strong reduction in EOP and frequently total elimination of mature plaques (fig. S6, S9-S19). For a few phages (T7, T5, EdH4, SUSP1, and M13), effective crRNAs primarily gave strong plaque size reduction, potentially indicative of imperfect repression of MCP.

**Fig. 4.**
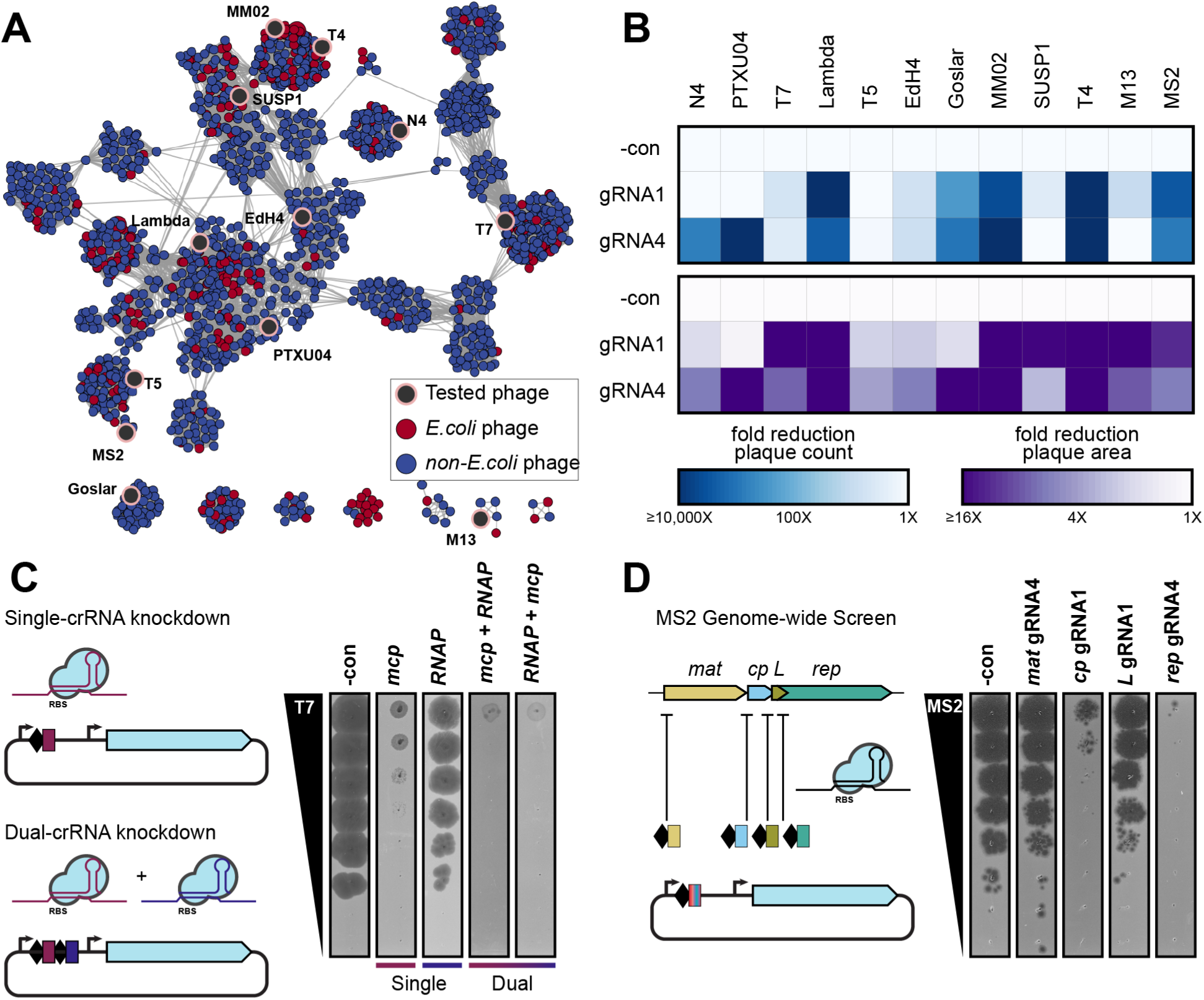
Translational repression is broadly active against phage diversity. (**A**)  Network graph representation of *E. coli* phages and their relatives (*23*). Nodes represent phage genomes connected by edges if they share similarity determined by vContact2 (*57*). Red and blue nodes represent *E. coli* and non-*E. coli* phages respectively. Phages assessed here for dCas13d sensitivity are shaded in black. (**B**) Anti-phage activity conferred by dCas13d when targeting an essential gene in 12 diverse phages (mean of three replicates), scored by EOP reduction (top) or plaque size (bottom). (**C**) crRNA multiplexing facilitates more efficient knockdown than component guides, using phage T7 as an example. (**D**) Genome-wide knockdown screen in phage MS2. Best of two guides tested are shown. dCas13d was induced as described in Methods.

CRISPRi-ART targeted against phage T7 showed a consistent reduction in plaque size but not EOP (Fig. 4B, S19). To test if plaque size reduction reflected an intermediate phenotype such as incomplete repression, we employed a dual crRNA targeting two T7 essential genes (T7*mcp* and T7*rnap*) alone or in combination (Fig. 4C). Both single crRNAs dramatically reduced plaque size without substantial EOP reduction. However, when targeting T7*mcp* and T7*rnap* simultaneously with two crRNAs, we observed nearly complete elimination of T7 plaque formation. These results suggest that CRISPRi-ART crRNA multiplexing can have a synergistic effect in the virocell.

We also found CRISPRi-ART to be effective against RNA phage MS2 (Fig. 4B). Because MS2 has a short, 3.6kb ssRNA genome (*41*), we extended CRISPRi-ART to target each of the 4 known CDSs in the MS2 genome: *mat* (maturase), *cp* (capsid), *L* (lysis protein), and *rep* (RNA-dependent RNA polymerase), all of which are essential (Fig. 4D). At least one crRNA for each MS2 gene inhibited infection, but crRNAs targeting outside of the susceptible RBS region on either +sense or -sense RNA strands did not, ruling out direct obstruction of genome synthesis (Fig. 4D, S14). We observed clear differences in magnitude of knockdown, with *rep* and *cp* being more sensitive to CRISPRi-ART than *L* or *mat*, potentially highlighting more important roles in MS2’s life cycle. In addition to highlighting CRISPRi-ART’s applicability to RNA phage biology, these results show the ability to perform genome-wide knockdown screens in phages. By leveraging the versatility of targeting RNA with dCas13d, CRISPRi-ART is capable of overcoming the genome-targeting obstacles imposed by diverse phages (Fig. 3A, 4).

## A pooled fitness screen enables genome-wide phage-gene essentiality assignment in diverse phages

Having demonstrated dCas13d’s ability to inhibit diverse phages with single crRNAs, we performed genome-wide CRISPRi-ART functional genomics screens (Fig. 5A) on a subset of the phages: model phages T4 (Fig. 5B) and T5 (Fig. 5C) as well as non-model phages SUSP1 (Fig. 5D) and PTXU04 (Fig. 5E). By directing dCas13d to genome-wide predicted phage RBSs in a pooled crRNA library, CRISPRi-ART enables the simultaneous interrogation of every gene’s role on phage infection (i.e., phage fitness). For the pooled fitness assays, we exposed the library supplemented with dCas13d and crRNA inducers, to phage at high multiplicity of infection (MOI - 10 or 100) on solid plate media (Methods). After 14 hrs of phage exposure, we recovered the surviving colonies for sample preparation and crRNA counts in surviving cells were compared to an uninfected control to quantify phage gene fitness scores (Methods). During these pooled CRISPRi-ART assays, crRNAs targeting RBS of an important gene essential for phage infection (for example, a phage capsid) confer a selective advantage to the host, increase in abundance, and therefore receive a positive fitness score. Here, a “Fit” phage gene means a specific CRISPRi-ART crRNA facilitated survival of the *E.coli* colony across 7 crRNAs (log2FC > variable threshold (Methods), p<0.05), suggesting that the targeted gene was essential or important for phage infection. “Semi-Fit” genes refer to genes with substantial crRNA variability (p > 0.05), but large fold-changes. Because this could reflect variability intrinsic to the crRNA and not the targeted gene content, we opted to separately denote this category.

**Fig. 5.**
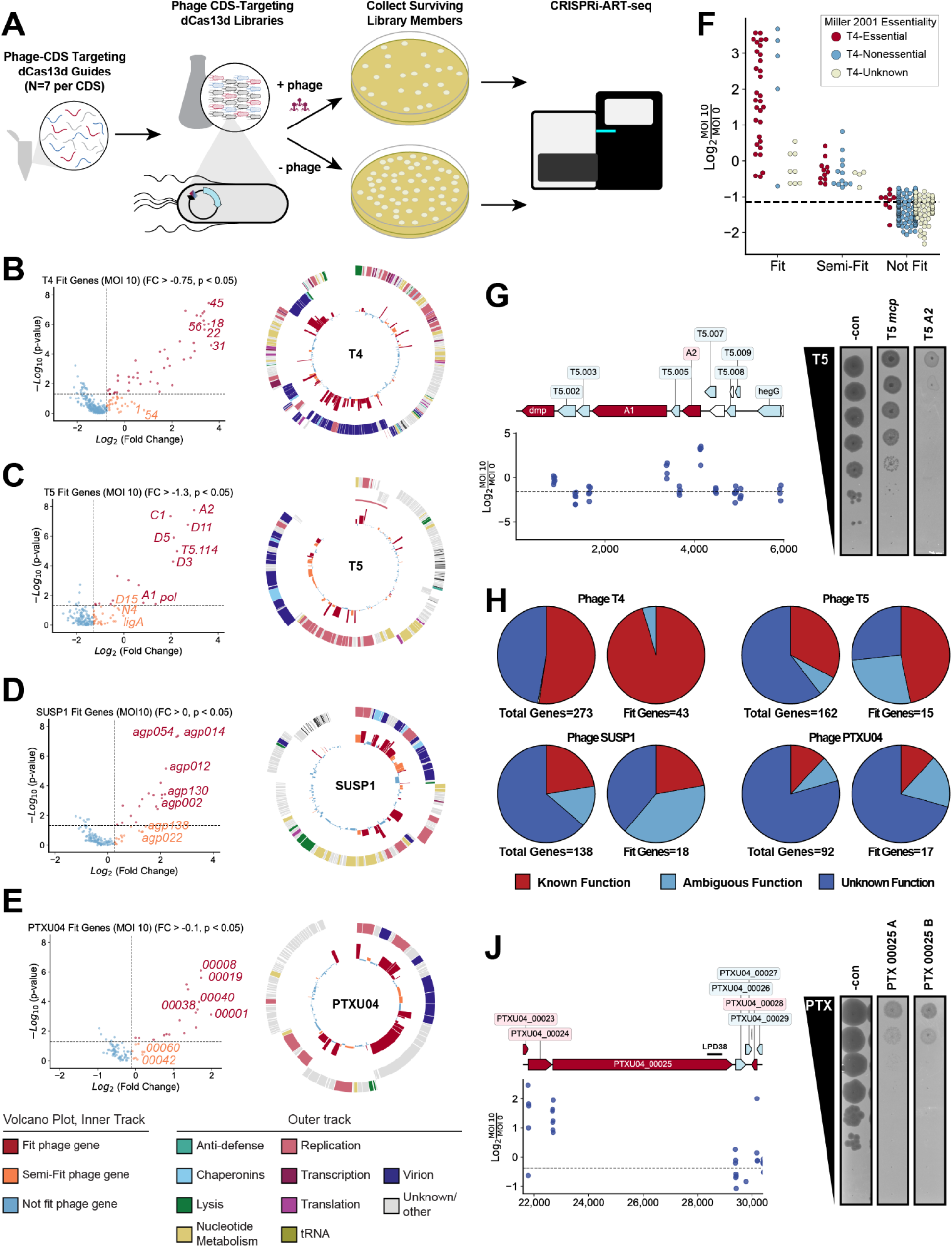
Programmable translational suppression unveils the gene essentiality landscape of phages. **(A)** Experimental library screening pipeline to discover phage genes important for efficient phage infection. (**B-E**) Volcano- (left) and Circos- (right) plots summarizing phage functional genomics screens for phages T4 (**B**), T5 (**C**), SUSP1 (**D**), and PTXU04 (**E**) Representative fit genes and semi-fit genes are highlighted in red and orange respectively. Thresholds for significance and fitness in Volcano plots are shown with a dashed line. In Circos plots, the outer track represents phage with broad functional categorization and the inner track displays fitness colored if it passes fitness criteria. (**F**) Distribution of fit and not fit genes in phage T4 colored by fitness from (B), binned by essentiality reported previously (*36*). (**G**) Phage crRNA fitness in pre-early genes in phage T5 (left) and plaque assays (right). (**H**) Distribution of phage gene-annotation quality for all phages across the full genome (left) and Fit genes (right). (**J**) Phage gene fitness in hypothetical, putative polyvalent protein encoded by *PTXU04_00025* (left) and plaque assays (right). Gene fitness is shown as the mean of 3 biological replicates with log10-transformed K-S p-value. crRNA fitness is shown as the median of 3 biological replicates.

Across all four genome-wide CRISPRi-ART phage screens, we generally observed strong enrichment of crRNAs targeting annotated structural phage genes (ex. *mcp*) (Fig. 5B-E), suggesting that positive fitness reflects an inability of a high MOI phage infection to efficiently produce viable progeny. An extended description of Fit phage genes across four phage assays is provided in supplementary text and put into full-genomic context in fig. S20-S23. To benchmark our genome-wide assays, we compared our T4 phage CRISPRi-ART data against known essential/non-essential gene roster of T4 phage (Fig. 5F) (*36*). Broadly, CRISPRi-ART was concordant with known assignment of gene essentiality in T4 phage, capturing 41 of 49 established T4 essential genes with fold-change alone. Our results also uncovered several additional Fit genes that are known to be essential in some hosts but not others. For instance, *dmd* encodes an antitoxin against a T4-induced toxin, RnlA, encoded in our *E. coli* assay strain (*42*). One gene, *49.1*, appeared as a Fit gene in our screens with no documented biological function, but highly conserved across T4-like phages, revealing previously unrecognized importance of this gene for T4 infection. An extended description of these results is provided in supplementary text.

Compared to phage T4, the top Fit genes uncovered in our CRISPRi-ART screen of T5 phage are known to play a role in phage replication cycle (ex. *D6* and *pol* encoding replicative DNA helicase and DNA polymerase respectively) and early infection step. (Fig. 5C). For example, genes important for first/second step transfer (FST/SST) (*A1* and *A2*), a unique feature of T5’s life cycle in which T5 injects ∼10kb of its genome in a discrete stage (FST) from the rest of its genome (SST) (*18*), were highly fit. To validate the relative importance of lifecycle coordination in phage T5 we investigated the impact of targeting gene *A2*, one of two uncharacterized essential genes involved in T5 SST DNA injection. *A2* was among the top fit genes in phage T5, along with essential gene *A1* and critical, albeit non-essential *dmp* (Fig. 5G) (*18*, *43*, *44*). Further inspection of a single crRNA targeting *A2* showed strong inhibition of phage T5 in comparison to *mcp*targeting crRNA (Fig. 5G, S28), reinforcing the relative importance of lifecycle control in phage T5.

## Genes of currently unknown function are central to phage infection

We next investigated Fit genes of unknown function within phage genomes (Methods). While T4 and T5 are among the most well studied phages, over 50% of their respective genes have no assigned role in the infection cycle (Fig. 5H). This trend is exacerbated in non-model phage SUSP1 and singleton phage PTXU04, with up to 75% of Fit genes having no assigned function (Fig. 5H). We identified 30 Fit genes of no known function: 2 in T4 (*49.1* and *5.3*), 6 in T5 (6 of 17 Fit genes), 9 in SUSP1 (9 of 19 Fit genes) and 12 in PTXU04 (12 of 17 Fit genes). Of particular note, for PTXU04 the majority of Fit genes had no inferrable function (Fig. 5H) (Methods), highlighting novel infection strategies encoded by this phage. An extended description of all putatively essential fit and semi-fit genes can be found in supplementary text. As a demonstration of CRISPRi-ART platform in assigning genome-wide gene essentiality roles, we validated our screen’s knockdown against PTXU04_00025, a 2,198 aa putative polyvalent protein with no confident annotations beyond the Large Polyvalent Domain 38 (LPD38) signature (Fig. 5J) (*45*). Such proteins are predicted to play complex roles in subverting host-defense during phage infection, but have not yet been shown to be essential for efficient phage infection (*45*, *46*). Single crRNA targeting of *PTXU04_00025* confirmed that the PTXU04_00025 protein is indeed critical for efficient infection (Fig. 5J).

## Discussion

Here we establish CRISPRi-ART as a foundational phage functional genomics technology that fundamentally redefines how diverse, core functions encoded by phages are identified and studied. CRISPRi-ART is able to knock down protein expression and elicit fitness phenotypes across phylogenetically diverse phages by targeting dCas13d to the RBS of phage or host transcripts. Owing to the scalable, multiplexable nature of crRNA, CRISPRi-ART can measure the genome-wide impact of protein expression knockdown. Leveraging this approach we report the first genome-wide essentiality maps of multiple diverse lytic phages (fig. S24-S27). In all cases, we corroborate known essential components of phage genomes, but also identify many genes of entirely unknown function core to efficient phage infection. Especially for non-model phages sharing low protein similarity with sequenced phages, such as PTXU04, this can be an exceptionally high number of genes, suggesting unique host takeover strategies compared to well-studied phages. Beyond basic understanding of phages, resolution of gene essentiality can allow for rapid inference of deletable gene content. In tandem with wild-type Cas13 phage gene editing strategies (*23*), CRISPRi-ART can guide phage genome reduction for phage therapy or microbiome editing applications (*4*, *47–49*). Furthermore, given the broad vulnerability of phages to CRISPRi-ART targeting independent of genome protection strategies (*15*, *17–19*, *36*, *39*), we anticipate CRISPRi-ART to enable phage functional genomics across phage diversity.

The ability to knock down protein expression of phage essential genes provides a global look into phage-encoded function *in situ*. For instance, alongside this study, CRISPRi-ART has proven critical to discover novel pre-nucleus compartmentalization and RNA export in *Chimalliviridae* (i.e. nucleus-forming phages) (*50*, *51*), whose enclosed genomes are broadly shielded from DNA targeting CRISPRi (*19*, *20*, *22*, *52*). Other individual crRNAs targeting essential genes such as those knocking down A2 from phage T5 or 00025 from PTXU04, can be simple experimental approaches to study enigmatic infection states of diverse prokaryotic viruses. As suggested by the Dmd knockdown phenotypes we observed with T4, CRISPRi-ART could be employed to discover phage-encoded inhibitors of native defense systems depending on the background strain context (*42*, *53–56*). CRISPRi-ART provides a versatile and scalable approach for discovery amidst the otherwise insurmountable complexity of phage-encoded genetic diversity.

## Acknowledgements

We thank members of the Cress lab, Doudna lab, Savage lab, Pogliano lab, Mutalik lab, the Innovative Genomics Institute, InCoGenTEC National Laboratory Project (Peter Otoupal and Kelly Williams) and m-CAFEs Science Focus Area for helpful discussions, encouragement, and feedback. The authors thank Tomas Hessler for an updated version of the phage network graph in Fig. 4A (*23*); Hitomi Asahara and Netra Krishnappa for help with sequencing; Harneet Rishi for advice on CRISPRi pooled fitness assays; and Adam Deutschbauer for advice on sequencing library preparation. Phage SUSP1 was a generous gift from S. Adhya. M.J.A thanks Elizabeth Hammond for their organization and encouragement. **Funding:** B.A.A. was supported by m-CAFEs Microbial Community Analysis & Functional Evaluation in Soils, a Science Focus Area led by Lawrence Berkeley National Laboratory based upon work supported by the US Department of Energy, Office of Science, Office of Biological & Environmental Research under contract number DE-AC02-05CH11231. M.J.A was supported by the National Science Foundation’s GRFP. J.R.P. was supported by funding from the Joint BioEnergy Institute (JBEI). E.A. and J.A.P. were funded by the Howard Hughes Medical Institute Emergent Pathogens Initiative grant. D.F.S. and J.A.D. are an Investigators of the Howard Hughes Medical Institute. B.F.C. was funded by U. S. Department of Energy, Office of Science, through the Genomic Science Program, Office of Biological and Environmental Research, under the Secure Biosystems Design project Intrinsic Control for Genome and Transcriptome Editing in Communities (InCoGenTEC); this work was also supported by a Research Award from the Shurl and Kay Curci Foundation (https://curcifoundation.org) to the Innovative Genomics Institute Genomic Tool Discovery Program at UC Berkeley, awarded to B.F.C.. **Author Contributions:** B.A.A., M.J.A., D.F.S., J.A.D., and B.F.C. conceived of the project. B.A.A., M.J.A., J.R.P., E.A., K.V.M., A.L., M.L.C., and B.F.C. performed experiments and built genetic constructs. B.A.A., M.J.A., J.R.P., E.A., and B.F.C. analyzed the data. E.C., V.K.M., J.S.S., and D.F.S. provided critical reagents and advice. B.A.A., M.J.A., and B.F.C. wrote the first draft of the manuscript. All authors contributed to writing and editing the manuscript. J.A.D. and B.F.C. supervised the project. **Competing interests:** The Regents of the University of California have patents pending related to this work on which B.A.A., M.J.A., J.R.P., E.A., E.J.C., J.A.P., D.F.S., J.A.D., and B.F.C. are inventors. J.A.D. is a co-founder of Caribou Biosciences, Editas Medicine, Intellia Therapeutics, Scribe Therapeutics and Mammoth Biosciences, a scientific advisory board member of Caribou Biosciences, Intellia Therapeutics, eFFECTOR Therapeutics, Scribe Therapeutics, Synthego, Mammoth Biosciences and Inari, and is a Director at Johnson & Johnson and has sponsored research projects by Biogen, Roche and Pfizer. V.K.M. is a co-founder of Felix Biotechnology. R.B. is a shareholder of Caribou Biosciences, Intellia Therapeutics, Locus Biosciences, Inari, TreeCo, and Ancilia Biosciences. The funders had no role in study design, data collection and analysis, decision to publish or preparation of the manuscript. **Data and materials availability:** Plasmids used in this study will be made available through Addgene following peer review of this manuscript. Illumina sequencing data used in this study will be deposited to the sequence read archive following peer review of this manuscript. Supplementary data for the preprint can be found on FigShare: https://doi.org/10.6084/m9.figshare.24154581.

