## Supplementary Information for "Genome-wide Characterization of Diverse Bacteriophages Enabled by RNA-Binding CRISPRi"

[1]California Institute for Quantitative Biosciences (QB3), University of California, Berkeley, California 94720, USA; [2]Innovative Genomics Institute, University of California, Berkeley, California 94720, USA; [3]Department of Molecular and Cell Biology, University of California, Berkeley, California 94720, USA; [4]School of Biological Sciences, University of California, San Diego, La Jolla, CA 92093, USA. [5]Graduate Group in Biophysics, University of California, Berkeley, Berkeley, CA 94720, USA; [6]Gladstone Institute of Data Science and Biotechnology, San Francisco, CA 94158, USA; [7]Department of Food, Bioprocessing and Nutrition Sciences, North Carolina State University, Raleigh, NC, USA; [8]Environmental Genomics and Systems Biology Division, Lawrence Berkeley National Laboratory, Berkeley, California 94720, USA; [9]Systems Biology Department, Sandia National Laboratories, Livermore, CA, USA 94550; [10]Howard Hughes Medical Institute, University of California, Berkeley, California 94720, USA; [11]Department of Chemistry, University of California, Berkeley, California 94720, USA; [12]MBIB Division, Lawrence Berkeley National Laboratory, Berkeley, California 94720, USA.

*Authors contributed equally.

### Materials and Methods

*Chemicals, reagents, and media*. All liquid, solid, and soft media were prepared with LB Broth Lennox (1% (w/v) tryptone, 0.5% (w/v) yeast extract, 0.5% (w/v) NaCl) and supplemented with antibiotics, inducers, and cations as needed and described below. Bottom and top agar were prepared with 1.5% (w/v) and 0.7% (w/v) agar, respectively. LB Broth Lennox was used for routine cultivation of *E. coli* and for experiments in rich medium. For experiments using alternative carbon sources, strains were cultured in M9 minimal medium (1X M9 salts, 2 mM $MgSO_4$, 0.1 mM $CaCl_2$) supplemented with 0.265 mg/mL thiamine HCl, 2 mg/mL casamino acids, and a primary carbon source (0.4% (v/v) glycerol, 20 mM D-galactose, 20 mM D-xylose, 20 mM D-maltose monohydrate, or 20 mM L-fucose). Expression of dCas13d was induced by the addition of anhydrotetracycline (aTc; Sigma-Aldrich, CAS 13803-65-1). crRNAs encoded on auxiliary crRNA expression plasmid pBFC1171 were induced using 1 μM crystal violet (for pooled crRNA libraries targeting *E. coli*), while crRNAs encoded on pBFC0984 were expressed from strong constitutive promoter BBa_J23119. Antibiotics were used at a concentration of 34 μg/mL for chloramphenicol and 100 μg/mL for carbenicillin. SM buffer (Teknova) was used for phage dilution. All bacterial and phage strains used in this work are listed in data S2, S3.

*Competent cell production*. Commercial chemically competent and electrocompetent cells were used when available. For custom chemical competent cells were cultured in ZymoBroth, and made competent using Mix & Go buffers (Zymo) according to the manufacturer's recommended protocol. For custom electrocompetent cells, Overnight *E. coli* cultures in 2xYT medium with appropriate antibiotics were inoculated into the same medium to an OD of 0.05 and grown to mid-exponential phase (OD600 0.4-0.6). Cells were pelleted, washed twice with chilled H2O and twice with chilled 10% glycerol, and resuspended in chilled 10% glycerol to achieve a ~300× concentration of the harvested culture. Aliquots were immediately frozen at -80°C.

*Full plasmid sequencing*. All plasmid constructs were sequenced by full-plasmid sequencing services at the UC Berkeley DNA Sequencing core (Illumina or Oxford Nanopore), Primordium (Oxford Nanopore), or Plasmidsaurus (Oxford Nanopore).

*Phage propagation and scaling*. Phages were propagated through commonly used protocols in LB media or LB top agar overlays (0.7%). Unless stated otherwise, phages were propagated on *E. coli* BW25113 (lacI+rrnBT14 ΔlacZWJ16 hsdR514

ΔaraBADAH33 ΔrhaBADLD78 rph-1 Δ(araB–D)567 Δ(rhaD–B)568 ΔlacZ4787(::rrnB-3) hsdR514 rph-1). Phages N4, T4, T5 and T7 were scaled on *E. coli* BW25113. Phage SUSP1 was a gift from Dr. Sankar Adhya and scaled on *E. coli* BW25113. Phages EdH4 and MM02 were obtained from the DSMZ culture collection and scaled on *E. coli* BW25113 (DSM 103295 and DSM 29475, respectively). Phage λ cI857 bor::kanR was a gift from Dr. Drew Endy and scaled as described previously. Phage MS2 was a gift from Vivek Mutalik and scaled in *E. coli* strain NEB 5-alpha F'Iq (NEB) with 2 mM $CaCl_2$. Phage M13 was obtained from ATCC (15669-B1) and was also propagated on NEB5α F'Iq genotype cells (NEB) with 2 mM $CaCl_2$. All phages were titered through 2 µl spots of 10× serial dilution of phage in SM buffer on *E. coli* BW25113 or NEB5α F'Iq in a 0.7% top agar overlay.

*CRISPRi-ART vector construction.* The *Ruminococcus flavefaciens* Cas13d (RfxCas13d) coding sequence was amplified from addgene plasmid pXR001: EF1a-CasRx-2A-EGFP, a gift from Patrick Hsu (Addgene plasmid # 109049 ; http://n2t.net/addgene:109049 ; RRID:Addgene_109049). The primary CRISPRi-ART vector pBFC0984 was constructed by assembling a p15A chloramphenicol resistant backbone with the catalytically deactivated dRfxCas13d coding sequence under transcriptional control of the aTc-inducible pTet promoter, along with a 2xBsaI Golden Gate spacer cloning site for expression of crRNAs from the strong constitutive promoter BBa_J23119. Plasmid pBA556 is similar to pBFC0984 but lacking the crRNA cassette. Plasmid pBFC0984 was used as a spacer entry vector for all individual and dual crRNA constructs, as well as for all phage crRNA libraries. In order to reduce likelihood of leaky crRNA expression that could lead to pooled crRNA library bias for libraries targeting the *E. coli* transcriptome, we also built a pBA556-compatible auxiliary crRNA vector pBFC1171 consisting of a low-copy SC101 origin, *bla* ampicillin/carbenicillin resistance marker, and a 2xBsaI Golden Gate spacer cloning site for expression of crRNAs from the non-leaky, strong, crystal violet-inducible promoter pJEx (*1*). Plasmid pBFC1171 was used as a spacer entry vector for all *E. coli* crRNA libraries. For control crRNA-only samples, plasmid pBFC0843 was constructed and used in place of pBA556 along with the pBFC1171-harbored crRNA library. Plasmid pBFC0843 is dRfxCas13d-null and crRNA-null and possesses a p15A origin, chloramphenicol resistance marker, and a pTet promoter without a downstream CDS.

*Individual and dual crRNA cloning*. To introduce crRNA spacers into pBFC0984, we employed BsaI-HFv2 (NEB, R3733L) Golden Gate assembly (*2*). For individual crRNAs, spacers were designed as two complementary oligonucleotides with 4 bp 5' overhangs that matched the staggered ends of the BsaI-digested destination plasmid. For dual crRNAs, two pairs of oligos (each encoding one of the two spacers and part of the central direct repeat) were designed to ligate into the backbone in a similar manner, inserting a spacer-repeat-spacer segment. These oligonucleotides were phosphorylated using T4 PNK (NEB) at 37 °C for 30 min and then duplexed at a concentration of 10 uM. Duplex formation involved melting at 100 °C for 5 min, followed by slow cooling to room temperature over a span of 15 min. The PNK-annealed spacer duplexes (100 fmol) served as the insert templates in each Golden Gate reaction. Cloning reactions were subsequently transformed into competent *E. coli* (Mach1-T1R, NEB10B, or IG10B) and clones verified by full plasmid sequencing.

*Quantifying CRISPRi-ART in E. coli using individual and dual crRNAs.* Plasmids containing a non-targeting control crRNA (the pBFC0984 2xBsaI non-targeting sequence), individual crRNAs targeting the RBS region of *gfp* or *rfp*, or dual crRNAs targeting *gfp*+*rfp* or *rfp*+*gfp* were transformed into *E. coli* MG1655 harboring genomically encoded RFP and GFP, as previously described (*3*). Plasmids containing a non-targeting control crRNA (our default RFP-targeting negative control plasmid) or individual crRNAs targeting the RBS region of *dnaK* were transformed into *E. coli* BW25113. Single colonies were picked into 5 mL Teknova EZ Rich Defined Media (20% glucose) supplemented with 34 µg/mL chloramphenicol and cultured overnight for 16hr to stationary phase in a 24 deep well plate using an Infors Multitron plate shaker (37°C and 800 rpm). Cultures were then diluted 1:1000 into the same medium supplemented with 200 nM aTc to induce dRfxCas13d expression. Induced cultures were collected in mid-log phase ($OD_{600}$ = 0.4) to quantify protein (western) and mRNA (qRT-PCR) abundance as described below. For RFP- and GFP-targeting experiments, 5 mL samples in mid-log phase were collected for mRNA quantification, and matched 5 mL cultures were allowed to continue growing to facilitate fluorescent protein maturation and measurement 16 hour post-induction. Endpoint fluorescence measurements were taken on a Biotek plate reader (RFP λex = 575 nm and λem = 610nm; GFP λex = 485 nm and λem = 535nm). Plate reader assays were performed at 200 µL in flat clear-bottom, black-walled 96-well plates with a non-binding surface treatment (Corning, catalog no. 237105).

Cells were centrifuged at 4000 g for 10 min at 4°C. Lysis was achieved by adding SoluLyse™ Bacterial Protein Extraction Reagent (Genlantis, L200125) according to the manufacturer's instructions. After incubation for 15 minutes at room temperature with gentle agitation, lysates were clarified by centrifugation at 12,000 x g for 20 minutes at 4°C. The supernatant, containing the soluble protein fraction, was collected for further analysis. 20–50 µg protein lysate was denatured in 1× Laemmli buffer at 95C for 10 min and resolved by SDS-PAGE. Protein was transferred to Immun-Blot LF PVDF membrane (Bio-Rad). The membrane was blocked with blocking buffer (PBS/0.05%

Tween-20 containing 5% milk) for 1 h at room temp, incubated with primary antibody in blocking buffer overnight at 4C, washed three times with PBS/0.05% Tween-20 for 5 min each, incubated with dye-conjugated secondary antibody in blocking buffer for 1 h at room temp and washed three times again with PBS/0.05% Tween-20 for 5 min each. Protein bands were visualized on an LI-COR Odyssey CLx with Image Studio v5.2 software using 700 nm and 800 nm channels. The following primary antibodies were used: mouse anti-DnaK (abcam, AB69617, 1037401-3, 1:2,000 dilution) and rabbit anti-GroEL (abcam, catalog AB90522, 1024635-2, 1:2,000 dilution); the following secondary antibodies were used: IRDye 680RD goat anti-mouse (LI-COR, 926-68070, Lot C90910-21, 1:20,000 dilution), IRDye 800CW goat anti-rabbit (LI-COR, 926-32211, Lot C90723-19, 1:20,000 dilution). The captured Western blot images were imported into Empiria Studio 3.0 software for quantification. Bands of interest were selected using the region of interest (ROI) tool. Background subtraction was performed using the rolling ball algorithm with a radius of 50 pixels. The integrated density of each band was measured and normalized to the corresponding loading control GroEL band. The data were then analyzed and plotted using GraphPad PRISM.

Total cellular RNA was extracted using TRIzol Reagent (Thermo Fisher Scientific) per manufacturer's instructions. Genomic DNA was removed using TURBO DNase (Thermo Fisher Scientific). After inactivating TURBO DNase with DNase Inactivation Reagent, 2 µg DNase-free RNA was reverse transcribed using SuperScript III Reverse Transcriptase (Thermo Fisher Scientific) with random primers (Promega) per manufacturer's instructions. qPCR was performed using iTaq Universal SYBR Green Supermix (Bio-Rad) in a CFX96 Real-Time PCR Detection System (Bio-Rad) using the following thermocycler conditions: 95°C for 3 min, [95°C for 15 s, 58°C for 30 s, 72°C for 30 s]x45 cycles. Gene-specific primer pairs used to detect transcripts are listed in data S4. Relative amount of a given target RNA under targeting versus non-targeting conditions was calculated using the formula 2^-((CtTarget–CtGroEL)Targeting crRNA – (CtTarget–CtGroEL)NT crRNA). Three biological and three technical replicates were run for each condition. PCR reactions were verified to produce a single band of expected size via agarose gel electrophoresis, and no-RT and no-template controls were run in parallel (not shown).

*Quantifying CRISPRi-ART against phages using plaque assays with individual and dual crRNAs.* Bacteriophage plaque assays were performed using a modified double agar overlay protocol as reported previously (*4*). Unless stated otherwise, phage assays were performed using DH10b-genotype *E. coli* (NEB, Intact Genomics) or DH5alpha F' genotype *E. coli* (NEB C2992) (for MS2 and M13 phages) transformed with a plasmid containing dRfxCas13d, dLbCas12a, or dSpyCas9 under pTet control with a crRNA (or sgRNA in the case of dSpyCas9) under constitutive control (data S1). Cultures were grown overnight at 37 °C and 250 r.p.m. with appropriate antibiotics, and 100 μl of saturated overnight culture was mixed with 5 mL molten LB Lennox top agar supplemented with appropriate inducer (below) and antibiotics. This mixture was decanted onto a corresponding 5 mL LB Lennox + chloramphenicol agar plate to final overlay concentrations of 0.7% (w/v) agar, aTc (variable, below) and 34 μg mL−1 chloramphenicol. For dCas13d experiments the following final concentrations of aTc were used to minimize background toxicity while maintaining phage inhibition: phages T4, MM02, and Lambda (20 nM), phage Goslar (50 nM), and phages EdH4, M13, MS2, N4, PTXU04, SUSP1, T5, and T7 (100 nM). For T4 phage experiments involving dLbCas12a a lower, final aTc concentration of 10 nM was used due to expression toxicity.

In general, no supplementary $CaCl_2$ or $MgSO_4$ salts were added except for experiments involving phages MS2 and M13, which employed a final concentration of 1 mM $CaCl_2$. Overlays were left to dry for 15 min under microbiological flame. For each Cas-crRNA-phage combination, 10× serial dilutions of the appropriate phage were performed in SM buffer (Teknova), and 2 μl of each dilution were spotted onto the top agar and allowed to dry for 10 min. Plaque assays were incubated at 37 °C for 12–16 hours. Post-incubation, plaques were scanned using a photo scanner and plaque forming units (PFUs) enumerated. When no plaques but clearings were observed at high phage concentrations, we considered these as “lysis from without” and indicated a lack of productive phage infection (*5*). We approximated these EOPs as 1 PFU at the most concentrated dilution of clearing. EOPs were calculated by normalizing the mean of PFUs for a condition to the mean PFU of a negative control (targeting RFP by default): mean(PFUcondition)/mean(PFUnegativecontrol). All plaque assays were performed in biological triplicate and EOP calculations performed using GraphPad PRISM.

Plaques were further analyzed by size in Fiji (*6*). Image scale was set to 0 and individual plaques were selected as regions of interest using the full plaque area including turbid zone. The area of each plaque was calculated. Fold-change for plaque size measurements was calculated as the following: mean(area_condition)/mean(area_control).

*Oligo pool design and amplification*. Oligo pools were synthesized by Twist Bioscience and were designed to encode PCR amplifiable crRNA libraries to be cloned into pBFC0984 or pBFC1171 using BsaI Golden Gate assembly. Oligos containing internal BsaI sites (due to BsaI in the target or a rare BsaI arising when concatenating the final oligo components) were excluded from synthesis to reduce assembly errors. Duplicate oligos encoding crRNAs targeting multi-copy or repetitive genomic features were deduplicated prior to synthesis. The crRNA libraries were synthesized as pools in which each distinct crRNA library was designed to be uniquely amplifiable with an orthogonal primer pair (*7*). Each oligo was composed of the following key sequence elements, concatenated in the 5' to 3' direction: a 20 nt subpool-specific forward primer, 11 nt encoding the upstream BsaI site and AAAC overhang, 31 nt variable spacer sequence, 1 nt to maintain the starting base of the downstream terminator feature on the crRNA entry vector, 11 nt encoding the downstream TGCT overhang and BsaI site, and a 20 nt subpool-specific reverse primer matched to the upstream primer, and 6 nt arbitrary DNA (fig. S2). Orthogonal primer pairs used for subpool amplification are listed in data S4). Sense RBS control oligos (described below) were synthesized as part of a separate oligo pool from antisense RBS-targeting oligos to prevent amplification problems arising from hybridization of these complementary oligos. Oligo pools were resuspended in Qiagen EB (10 mM Tris, pH 8.5) to 10 ng/uL and stored at -80°C when not in use. Subpools were amplified using subpool-specific primers following Twist recommendations and the KAPA HiFi HotStart DNA Polymerase Kit (Material Number: 7958889001). Specifically, 25 µL reactions were assembled with 0.5 U KAPA HiFi HotStart DNA Polymerase, KAPA HiFi Fidelity Buffer, 0.3 mM dNTPs, 5 ng oligo pool, 0.3 µM each subpool-specific primer. The thermocycler program included an initial melting step for 3 min at 95°C; 8 cycles of 98°C melting for 20 s, 50°C annealing for 15 s, 72°C extension for 15 s; and a final extension at 72°C for 1 min. Bioanalyzer confirmed successful amplification of the expected 98 bp products. These PCR products were purified using a DNA Clean & Concentrator-5 kit (Zymo, D4004), eluted with 10 µL milliQ $H2O$, and used to assemble crRNA libraries as described below.

*E. coli CRISPRi-ART single nucleotide-tiling crRNA library design*. We designed a pooled, single nucleotide resolution library with 29,473 crRNAs tiled anti-sense to 18 *E. coli* BW25113 (Accession CP009273) transcripts, 16 containing at least one essential gene and 3 containing a counterselectable gene (data S5). crRNAs were tiled 100 nt beyond the ends of transcriptional start and stop sites when known (*8*), or 100 nt beyond outermost coding sequences when not previously reported. Metadata for the single nucleotide-tiling crRNA library is shown in data S5.

*E. coli CRISPRi-ART RBS-targeting crRNA library design*. The *E. coli* genome-wide crRNA library was designed to target the RBS region of all known and hypothetical CDSs in *E. coli* BW25113, using 7 crRNAs coarsely tiled in 5 nt increments antisense to the susceptible RBS region highlighted in Fig. 1E (fig. S1), totaling 31,902 crRNAs. This enabled comprehensive genome-wide coverage, with at least one guide overlapping both the RBS and start codon of the target gene. A separate sense (non-targeting) control oligo library was designed to contain 1 crRNA for each gene, identical to the coding strand and spanning the RBS and start codon, totaling 4,571 crRNAs. These sense oligos were not anticipated to possess a target transcript and were therefore not expected to produce any fitness effects.

*E. coli CRISPRi-ART pooled crRNA library construction*. Given the higher diversity of our *E. coli*-targeting libraries, we used a crRNA library construction approach aimed at maintaining high library coverage and avoiding bias in the cloning and propagation steps. PCR products containing the crRNA libraries were cloned into pBFC1171 using BsaI Golden Gate assembly. To remove undigested entry vector, reactions were subsequently treated with a followup digestion and cleanup procedure, consisting of BsaI digestion at 37°C for 1 hr, BsaI heat inactivation at 85°C for 20 min, Plasmid-Safe ATP-Dependent DNase (LGC Biosearch Technologies) exonuclease treatment at 37°C for 1 hr, PlasmidSafe heat inactivation at 75°C for 30 min, and purification using a DNA Clean & Concentrator-5 with 10 uL elution in milliQ $H_2O$. High competency Endura (LGC Biosearch Technologies) electrocompetent cells were electroporated with 1 µL of this product and recovered at 37°C and 250 r.p.m. for 1 hr. A small aliquot was serially diluted and spot plated to count colonies, to estimate library coverage, and to sequence 10 colonies to confirm efficient and diverse crRNA insertion, and the remainder of the recovery stored at 4°C overnight. Based on transformation titers, an appropriate volume of the recovery was plated onto pre-dried bioassay dishes containing LB agar plus carbenicillin, aiming for 100× cfu over library size and no more than 1M cfu on a single bioassay dish. After 14 h overnight growth at 37°C, colonies were scraped from each bioassay dish into 50 mL LB plus carbenicillin, vortexed thoroughly, pooled if spread across multiple dishes, pelleted by centrifugation, and midiprepped with 200 uL milliQ $H_2O$ elution (ZymoPURE II Plasmid Midiprep Kit, D4201) to harvest plasmid library DNA (data S1, S2). To ensure complete removal of undigested entry vector from the plasmid library, 2 µg of DNA was treated with the followup digestion and cleanup procedure described above.

Next, experimental strain *E. coli* BW25113 was transformed with either pBA556 to build strain sBFC0264 or pBFC0843 to build strain sBFC0265 and subsequently made electrocompetent in preparation for transformation of crRNA library DNA. For the *E. coli* genome-wide RBS-targeting library, antisense and sense crRNA libraries were combined in a 7:1 molar ratio before transformation to account for 7 antisense crRNAs for each control sense crRNA (36,473 total). Both the single nucleotide tiling library and the pooled RBS-targeting libraries were electroporated into sBFC0264 and sBFC0265 (crRNA-only control library) and recovered at 37°C 250 r.p.m. for 1 hr. A small aliquot of the recoveries were serially diluted and spot plated onto LB agar plus chloramphenicol and carbenicillin to count colonies, to estimate library coverage, and to sequence 10 colonies to confirm maintenance of diverse crRNAs. The remainder of the recoveries were inoculated into 20 mL pre-warmed LB plus chloramphenicol and carbenicillin, grown at 37 C 250 r.p.m. until OD600 = 0.4-0.8, mixed with equal volume 40% sterile glycerol, and frozen at -80°C as 200 µL 20% revivable glycerol stocks. One glycerol stock for each library was thawed and titered on selective LB agar, indicating high viability after thawing with sufficient cfu to maintain high library coverage.

*E. coli pooled competitive fitness assays*. Library aliquots (200 µL) were thawed on ice for 10 minutes. One aliquot for each library was inoculated into 3 mL LB plus chloramphenicol and carbenicillin in a deep 24 well block and cultivated in a Multitron (Infors) plate shaker at 37°C and 750 r.p.m. until OD600 = 0.5-1.0. At this point, each culture was centrifuged at 4000 rcf for 5 minutes, supernatants aspirated, and pellets gently washed in 1 mL M9 base medium (without casamino acids and without a carbon source). This wash procedure was repeated for a total of three times before a final resuspension in 3 mL M9 base medium, and 30 µL of well mixed cells were inoculated into 3 mL of fresh assay medium, aiming for an initial cell count of 3E7 cfu. All assay medium contained the relevant base medium, antibiotics, and 200 nM aTc for dRfxCas13d induction and 1 µM crystal violet for crRNA induction. LB Lennox was used as the base assay rich medium for single nucleotide-tiling (Fig. 1D-E) and genome-wide RBS-targeting libraries (Fig. 2C) were grown in LB. M9 medium supplemented with the various carbon sources noted above was also used as base assay media for genome-wide RBS-targeting libraries (Fig. 2D). Competitive growth assays proceeded in 24 deep well blocks at 37°C and 750 r.p.m. until OD600 = 0.5-1.0 (7-8 doublings), at which point 30 µL of well mixed culture was subcultured into the same fresh assay medium and the remainder of the culture pelleted and frozen for subsequent CRISPRi-ART-seq of the intermediate time point. The final cultures were cultivated under the same conditions for another 7-8 doublings before harvesting and freezing at -80°C for subsequent CRISPRi-ART-seq, totalling 14-16 doublings post-induction.

*Phage CRISPRi-ART crRNA library design*. The genome-wide phage-targeting libraries were designed in the same manner as the *E. coli* coarsely-tiled antisense RBS library described above, including 7 antisense crRNAs (fig. S1) for each CDS. For each phage used in this study, gene coordinates and start codon annotations were obtained directly from NCBI, using the accession numbers phage T4 (NC_000866), T5 (NC_005859), SUSP1 (NC_028808), and PTXU04 (NC_048193).

*Phage CRISPRi-ART pooled crRNA library construction*. Given the lower diversity of our phage-targeting CRISPRi-ART libraries, we used a simpler approach for crRNA cloning in which our libraries were transformed into NEB10Beta (the assay strain for T4 and SUSP1, and the cloning strain for T5 and PTXU04). Oligo amplification, Golden Gate assembly, and followup digestion and cleanup steps were performed as described above, except pBFC0984 was used as the entry vector. Commercial electrocompetent NEB10Beta cells were then transformed with 1 µL of plasmid library DNA and recovered at 37°C 250 r.p.m. for 1 hr. A small aliquot of each recovery was serially diluted and spot plated on LB agar plus chloramphenicol to titer the transformations, and the remainder of the recoveries were stored at 4°C overnight. Transformation efficiencies were high, producing at least 100× greater cfu than library size, and sequencing of 10 colonies confirmed high cloning efficiency and crRNA diversity. Based on cfu count, an appropriate volume of recovery was plated on standard pre-dried LB agar plus chloramphenicol plates to obtain 100× cfu over library size, aiming for less than 100k cfu per plate. After 14 h overnight growth at 37°C, colonies were scraped from each plate into 50 mL LB plus chloramphenicol, vortexed thoroughly, and cultivated in non-baffled shake flasks at 37°C and 250 r.p.m.. After 3 hr cultivation, 8 mL of culture was mixed with an equal volume 40% sterile glycerol and frozen at -80°C as 1 mL 20% revivable glycerol stocks. The remainder of the culture was pelleted and midiprepped. The T5 and PTXU04 plasmid library DNA samples were further treated with follow up digestion and cleanup steps and then electroporated into their final assay strains *E. coli* IG10Beta and BL21, respectively, and validated and stocked as described here for NEB10Beta.

*Phage pooled crRNA competitive fitness assays*. Library aliquots (1 mL) were thawed on ice for 10 min, and then inoculated into 25 mL LB Lennox plus chloramphenicol in non-baffled flasks and cultivated at 37°C and 220 r.p.m. for 30 min. At this point the cultures were adjusted to 200 nM aTc to induce dRfxCas13d expression. Cultures were grown under induction at 37°C and 220 r.p.m. for another 1.5-2 hr. $OD_{600}$ measurements were used to estimate bacterial cfu/mL, and approximately 5E5 cfu (to ensure at least 100× coverage over library size) were set aside for plating. To these tubes containing *E. coli* expressing CRISPRi-ART libraries, phage stocks diluted in SM buffer were added to an MOI of 0, 10, or 100 mixed, allowed to adsorb for 15 minutes, and then plated on pre-dried LB agar plus chloramphenicol plates containing 200 nM aTc. After overnight incubation at 37°C, all colonies from a given plate were pooled into 10 mL LB, pelleted, and frozen at -80°C for subsequent CRISPRi-ART-seq.

*CRISPRi-ART-seq*. The frozen samples from the fitness experiments were thawed at room temperature after being stored at -80°C. The CRISPRi-ART crRNA library was isolated using a QIAprep® Spin Miniprep Kit. A PCR reaction was performed using 10ng of DNA from each sample in a 25 uL reaction volume, utilizing Q5 Hot Start polymerase and two primers (ctacacgacgctcttccgatctnnnnnctaccaactggtcggggtttg) and (cagacgtgtgctcttccgatctnnnnncctcttctgagatgagtttttgttcg) for P1 appending and downstream index appending. The P1 appending reaction consisted of an initial denaturation at 98°C for 30 seconds, followed by 15 cycles of denaturation at 98°C for 10 seconds, annealing at 55°C for 15 seconds, extension at 72°C for 10 seconds, and a final extension at 72°C for 2 minutes. The PCR reactions were then subjected to size selection using SPRISELECT beads according to the manufacturer's instructions, targeting product sizes of 250 bp. The eluted samples were used in a subsequent 50 uL PCR reaction with index primers (NEBNext® Multiplex Oligos for Illumina® (Dual Index Primers Set 2; Catalog No. E7780S)) for index appending. The index appending reaction involved an initial denaturation at 98°C for 30 seconds, followed by 5 cycles of denaturation at 98°C for 10 seconds, annealing at 55°C for 15 seconds, extension at 72°C for 10 seconds, and a final extension at 72°C for 2 minutes. Indexed samples were purified using SPRISELECT beads with the same size selection procedure. The purified samples were eluted in 20 uL of MilliQ water. Quantification of the purified samples was performed using a KAPA Library Quantification Kit (KAPABIOSYSTEMS, KR0405-v8.17), and the size of the products was confirmed using a Bioanalyzer 2100 automated electrophoresis system with a DNA 1000 Kit (Agilent Technologies, 50671504). The final samples were then sequenced on either an Illumina iSeq, Miseq, or pooled on a NextSeq, as specified in (data S6).

*crRNA read counting, normalization, and fitness calculations*. crRNAs were counted with 2fast2q (*9*, *10*) (https://github.com/afombravo/2FAST2Q) using the command python3 -m fast2q -c --m 0 --st 30 --l 31 --ph 0 and an input .csv file containing crRNA ID and corresponding spacer sequence for all crRNAs within the counted crRNA library. To account for variations in read depth between individual samples, raw feature counts were internally normalized for each tested crRNA by converting to reads-per-million. To account for denominator effects, 1 was added to each raw feature count prior to this normalization. For *E. coli* crRNA logfold-change (FC) calculations, crRNA feature counts in test samples were averaged across replicates and divided by corresponding counts in the T=0 condition. 2-way t statistic p-values were calculated using the scipy.stats module in Python3. FC underwent a log base 2 transformation using the numpy module in Python3. For phage FC calculations, the smallest and largest decile of crRNA (by read_counts) for MOI 0, 10, or 100 conditions were discarded to eliminate extrema prior to plating from gene fitness calculations. Remaining crRNA feature counts in test samples were averaged across replicates, and divided by corresponding counts in the MOI = 0 condition. To determine whether a gene conferred positive or negative fitness to a phage, the distribution of crRNA FCs for each gene was compared to the distribution of crRNA FCs across the entire phage genome. A bidirectional Kolmogorov–Smirnov test (K-S test) (via the scipy.stats module in Python3) was used to establish whether these distributions significantly differed.

*Determination of significantly impacted genes.* Related to Fig. 2D and fig. S3. To identify genes with fitness impacts in indicated conditions, we defined "significantly impacted" genes as those where at least 3 out of 7 tested crRNAs conferred a $\log_2$(FC) below -2, with all calculations performed with numpy and scipy.stats modules in python3. In Fig. 2D, the $\log_2$(FC) and t statistic p-values of the top 3 most differentially abundant crRNAs are averaged for each gene and plotted.

*Phage genome annotation*. T4, T5, SUSP1, and PTXU04 bacteriophage genomes were functionally annotated through a combination of automated and manual methods. T4, T5, and SUSP1 phages were automatically annotated using genomic annotations from CD-SEARCH (*11*) and PHROGS (*12*). Because PTXU04 CDSs exhibited limited relation to known proteins and PTXU04 is not in the PHROGS database, PTXU04 was manually annotated with hhpred on MPI Bioinformatics Toolkit (using PDB_mmCIF70_18_Jun, COG_KOG_v1.0, NCBI_Conserved_Domains(CD)_v3.19, and PHROGs_v4 domain databases) (*13*). Each PTXU04 gene was manually assigned an annotation based off of either a clear, confident hit (E-value <1e-5) or repeated low-confidence annotations (eg. phage tail protein). Additional attempts to annotate remaining hypothetical PTXU04 genes were performed using alphafold prediction followed by structural alignment to PDB and AFDB to limited success (*14–17*). All annotations were further manually inspected against known gene content in model phages T4 (*18*), model phage T5 (*19*, *20*), non-model phage SUSP1 (via progressiveMauve alignment with default parameters to related *Salmonella* phage Felix O1) (*21*, *22*), and non-model phage PTXU04 (*23*). Additionally, phage genomes were annotated with phage-defense inhibitors found in T4 and T5 phages (*24–27*). If conflict arose during annotation assignment, annotations were prioritized with the following confidence heuristic: literature > PHROGs > CD-SEARCH > hhpred. Any deviations were made based on annotation detail and annotation confidence.

In addition to the above annotations, genes were assigned "class" and "Annotation quality" scores. The "class" annotation included the following annotations: anti-defense, chaperonin, lysis, nucleotide metabolism, replication, transcription, translation, tRNA, virion, or unknown/other. "anti-defense" refer to genes involved in subverting phage defense systems including restriction modification systems. "chaperonin" refers to genes involved in phage virion or protein maturation, but not a structural component of the phage virion. “lysis” refers to genes involved in lysis, regulation of lysis timing, or degradation of cell wall components. “nucleotide metabolism” refers to genes responsible for nucleotide biosynthesis, degradation, modification, and regulation thereof, but not directly a part of replication. “replication” refers to genes involved in phage replication liberally applied. “transcription” refers to genes that modulate transcription in either the phage or host genome. “translation” refers to genes that modulate translation including genes that modulate RNA or tRNA stability. “tRNA” refers to tRNA genes, but not genes that modify them. “virion” refers to genes that are structural components or part of the virion produced in infection. “unknown/other” refers to all other genes encoded in phage. “Annotation quality” was assigned manually based off of both confidence, detail, and known literature of the annotation and its source content: “Known” for genes with known function, “Ambiguous” for known genes with ambiguity to substrate or role of the gene, and “Unknown” for genes of unknown function.

In general, these were in agreement with PHROG category with the following exceptions for visualization simplicity: (1) all predicted phage structural components were grouped into the “virion” category included packaged phage proteins, (2) many genes that are critical for phage lifecycle, but of unknown molecular function (ex. T5 genes *A1* and *A2*) were grouped into “replication”, (3) any gene responsible for assisting folding or assembly was overridden to fall under the “chaperonins” category, (4) genes responsible for anti-phage-defense through nucleotide modification were labeled as “nucleotide metabolism”, (5) genes with overlapping category functions (ex. RNase H were labeled with a primary annotation based on literature(*18*)), and (6) predicted subgenomic mobile elements (ex. homing endonucleases) were assigned “unknown/other” for simplicity. All phage annotations are listed in data S6).

*Analysis of phage CRISPRi-ART-seq*. Following CRISPRi-ART-seq processing, phage genes were interpreted for fitness. To identify FC thresholds for Fit and Semi-Fit genes, for each phage-MOI condition, we identified the lowest fitness score with a K-S p-value less than 0.05 on the right tail of the fitness distribution (ie positively enriched). This value was used as an inclusive FC threshold for fitness. Thus, we determined Fit genes through the following metrics: T4 (10 MOI) ($FC > -0.75$, $p < 0.05$), T4 (100 MOI) ($FC > -2$, $p < 0.05$), T5 (10 MOI) ($FC > -1.3$, $p < 0.05$), T5 (100 MOI) ($FC > -3.5$, $p < 0.05$), SUSP1 (10 MOI) ($FC > 0.25$, $p < 0.05$), SUSP1 (100 MOI) ($FC > 0$, $p < 0.05$), PTXU04 (10 MOI) ($FC > -0.1$, $p < 0.05$), PTXU04 (100 MOI) ($FC > -0.5$, $p < 0.05$). Genes passing FC thresholds, but not statistical thresholds were considered to be Semi-Fit; such cases are likely reflective of crRNA variability and typically reflected strong fitness of more than one crRNA per gene, but were underpowered to call Fit with significance. Due to the strong selection pressure baseline imposed by phage predation at high MOI we assumed that the average library member was reduced in abundance relative to MOI 0. Thus, we refrained from interpreting significant negative fitness scores from phage assays in this study.

Volcano plot visualization of phage gene fitness was visualized using python in matplotlib using FC for the mean of 3 biological replicates and -log10-transformed p-values. Circos plots were generated in python using pycircos (github.com/ponnhide/pyCircos) using phage annotations and mean fitness scores (and Fit/Semi-Fit/Not Fit classifications) from CRISPRi-ART-seq as outer and inner tracks, respectively. Genome-wide and genomic-region visualizations were generated using dna_features_viewer (https://edinburgh-genome-foundry.github.io/DnaFeaturesViewer/) and matplotlib in python. Coloring of phage genes was assigned by CRISPRi-ART-seq fitness classification and gRNA fitness displayed as the median of 3 biological replicates. Median gRNA fitness across the entire phage genome was shown with a dashed line. Comparison of T4 gene fitness scores to T4-essentiality (*18*) was performed using gene fitness scores and Fit, Semi-Fit, and No Fit classifications for the MOI 10 T4 infection condition using seaborn in python.

Guide RNA effectiveness targeting phage genomes (fig. S24-S27) was performed by analyzing the crRNAs targeting the top 10 fit genes in phages T4, T5, SUSP1, and PTXU04 (i.e. targeting genes that are clearly Fit). To assess absolute and relative gRNA effectiveness, the distribution of crRNA fitness scores and within-gene zscore (scipy.stats), respectively, were plotted using seaborn in python. Additionally, guides were interpreted by rank and the number of Top 3 guide RNAs within a gene were plotted by gRNA number using seaborn in python.

Supplementary Text:

Supplementary Text S1. *Knockdown via translational repression and or mRNA destabilization.*

Our data in supplementary figure S5 suggests that CRISPRi-ART may act through a combination of translational repression and mRNA destabilization. What is striking is in the case of DnaK gRNA1, we observe an mRNA level that is unchanged relative to the control and yet we observe an 82.7% knockdown in protein abundance. Suggesting that the majority of knockdown is due to translational repression rather than mRNA destabilization. This is then further implicated by the mRNA abundance of DnaJ, the downstream gene in the same transcriptional unit being relatively unchanged.

Supplementary text S2. *Summary of Genome-wide screens in T4.*

Our screens identified 45 Fit genes in model phage T4: *3*, *5.3*, *6*, *7*, *8*, *9*, *11*, *12*, *13*, *15*, *16*, *18*, *19*, *20*, *21*, *22*, *23*, *24*, *26*, *31*, *32*, *34*, *35*, *36*, *37*, *38*, *39*, *42*, *45*, *47*, *48*, *49.1*, *51*, *52*, *55*, *56*, *67*, *57A*, *dmd*, *goF*, *rnh*, *rpbA*, *t*, *vs*, and *uvsX*. Phage T4 is a large dsDNA phage with systematic efforts to fully annotate its genome (*18*). These efforts, while still incomplete - approximately one third of its genes remain entirely of hypothetical function - make T4 an ideal model phage to benchmark success of CRISPRi-ART in phage genomes.

Of the 45 Fit genes in model phage T4, we 43 had clear annotations: *3* (tail tube), *6* (baseplate wedge subunit), *7* (baseplate wedge subunit), *8* (baseplate wedge subunit), *9* (baseplate wedge tail fiber protein connector), *11* (baseplate wedge subunit), *12* (short tail fibers), *13* (neck protein), *15* (tail sheath stabilizer), *16* (small terminase subunit), *18* (tail sheath), *19* (tail tube), *20* (portal vertex), *21* (head maturation protease), *22* (head scaffolding protein, *23* (major capsid protein), *24* (capsid vertex), *26* (baseplate hub), *31* (head assembly chaperone), *32* (single stranded DNA binding protein), *34* (long tail fiber), *35* (tail fiber hinge connector), *36* (tail fiber hinge connector), *37* (long tail fiber), *38* (tail fiber assembly chaperone), *39* (DNA topoisomerase II large subunit), *42* (dCMP hydroxymethylase), *45* (sliding clamp), *47* (endonuclease subunit), *48* (tail tube junction protein), *51* (baseplate hub), *52* (DNA topoisomerase II medium subunit), *55* (late transcription sigma factor), *56* (dCTP pyrophosphatase), *67* (prohead core protein), *57A* (tail fiber chaperone), *dmd* (RnlA anti-toxin), *goF* (mRNA metabolism modulator), *rnh* (RnaseH), *rpbA* (RNA polymerase binding protein), *t* (holin), *vs* (valyl tRNA synthetase modifier), and *uvsX* (UvsX RecA-like recombination protein). An additional 2 Fit genes have no known function: *5.3* and *49.1*.

In addition we observed 26 Semi-Fit genes in T4, 18 with clear annotations: *1* (deoxynucleoside monophosphate kinase), *2* (DNA end protector protein), *4* (head completion protein), *10* (baseplate wedge subunit), *14* (neck protein), *25* (baseplate wedge subunit), *27* (baseplate hub), *29* (baseplate hub and tail length determinator protein), *41* (DnaB Replicative helicase), *43* (DNA polymerase), *44* (clamp loader), *54* (tail tube), *60* (DNA topoisomerase II), *imm* (superinfection immunity protein), *mobB* (homing endonuclease), *rI* (lysis inhibition regulator), *rIII* (lysis inhibition regulator), and *rnlA* (RNA ligase A). In addition, we observed 7 T4 Semi-Fit genes with purely unknown function: *5.4, 30.1*, *a-gt.4*, *cd.5*, *motA.1*, *trna.2, uvsY.-2*, *vs.1*, and *vs.8*.

While much has been learned about the T4 genome since then, (*18*) remains the most comprehensive, yet consolidated resource on the T4 genome and the encoded functions within. One particularly useful resource provided in this work is a careful accounting of empirical essentiality and non-essentiality of genes in the T4 genome, estimating a total of 49 individually essential genes in the T4 genome. Benchmarked against this resource, we recall 31 of these documented essential genes with high confidence and 43 including Semi-Fit genes. Our screens fail to capture fitness for 6 of these genes: 5 (tail lysozyme), 17 (Terminase large subunit), 33 (late promoter transcriptional regulator), 62 (clamp loader), 68 (prohead core protein), and e (endolysin). One insight we drew from this analysis was a limitation of CRISPRi-ART: multiple start sites for the same protein. T4's Large Terminase encoded by gene *17* is encoded by multiple start sites (fig. S20) (*28*). Given our model of function, we believe that dCas13d is occluding only one RBS at a time and potentially the Large Terminase is being produced in alternative isoforms from the same transcript. Future work is needed to determine if this is indeed the case. Nonetheless, CRISPRi-ART captures a very complete picture of essentiality in phage T4 when compared to deep literature review.

Our screens capture an additional 14 Fit and 15 Semi-Fit genes beyond what is listed in literature as T4-essential. Some of these genes are ostensibly essential due to clear ties to replication or virion structural components, but not proven empirically: *16* (small terminase subunit), *21* (head maturation protease), *29* (baseplate hub and tail length determinator protein), and *48* (tail tube junction protein). Some of these genes reflect genes important, but are documented not strictly essential for phage infection: 31 (GroES Chaperone), 32 (single-stranded DNA-binding protein), Rnh (RNase H), 47 (host-genome phosphoesterase), vs (Vs valyl-tRNA synthetase modifier), RpbA (RNA polymerase binding protein), GoF (mRNA metabolism modulator), *rnlA* (RNA ligase A), and *uvsX* (UvsX RecA-like recombination protein). An additional few may reflect a true population-level decrease in fitness, but not essentiality due to effects on suboptimal lysis timing (*rI* and *rIII*) or superinfection exclusion (*imm*). One homing endonuclease, MobB, showed up as fit that is likely fully dispensable.

One particularly interesting result was T4-encoded *dmd* emerging as a Fit gene. Dmd encodes for an antitoxin inhibiting *E.coli* -encoded toxin RnlA, whose activity otherwise degrades RNA. Inhibition of RnlA by Dmd stabilizes T4 mid-infection, conferring a strong benefit by Dmd (*29*). While Dmd is non-essential in the context of RnlA- infection, DH10b harbors an intact copy of RnlA, suggesting that CRISPRi-ART screens can identify defense system inhibitors encoded in lytic phage genomes. Notably anti-defense genes that were not relevant to the strain used in this screen do not appear phage fit (*24–26*).

Finally, we highlight the inclusion of 11 genes of hypothetical function that appear Fit (*5.3* and *49.1*) or Semi-Fit (*5.4*, *30.1*, *a-gt.4*, *cd.5*, *motA.1*, *trna.2*, *uvsY.-2*, *vs.1*, and *vs.8*). These genes represent potential inroads into new, previously unidentified biology encoded by phage T4.

Supplementary Text S3. *Summary of Genome-wide screens in T5.*

Our screens identified 17 Fit genes in model phage T5: *dmp*, *A1*, *A2*, *C1*, *T5.047*, *T5.084*, *T5.089*, *T5.090*, *D3*, *T5.114*, *D5*, *D6*, *pol*, *D11*, *T5.148*, *D20-21*, and *oad*. While phage T5 is among the original model type phages (*30*), it remains comparatively uncharacterized relative to model phages such as T4. While key features of T5 genetics remain a mystery, mechanisms in phage-defense and phage-defense-subversion mechanisms have renewed interest in this phage.

Of the 17 Fit genes in model phage T5, 9 had clear, known annotations: *dmp* (5′-Deoxyribonucleotidase), *A1* (SST DNA injection essential protein), *A2* (SST DNA injection essential protein), *C1* (holin), *D6* (replicative DNA helicase), *pol* (DNA polymerase), *D11* (putative single strand DNA binding protein), *D20-21* (major capsid protein), and *oad* (long tail fiber). Additionally 2 putative DNA-binding regulatory proteins, *D3* and *D5* appeared Fit as well. The remaining 6 Fit genes (*T5.047*, *T5.084*, *T5.089*, *T5.090*, *T5.114*, *T5.148*) are of entirely unknown function.

One of the most striking results when screening across the T5 genome is the clear enrichment of crRNAs targeting pre-early genes: dmp, A1, and A2. Following phage adsorption to the cell, T5 injects the first 10kb of its genome (encompassing 17 genes) into the cytoplasm, beginning a phase called first step transfer (FST) (*20*). Following expression and activity of A1 and A2 (the only strictly essential genes in this region), T5 injects the remainder of its genome by an unknown mechanism in a phase called second step transfer (SST). Our results corroborate the relative importance of A1 and A2 activity. Additionally host-genome degrading protein Dmp, shows fitness in our screens, deviating from established non-essentiality of this gene (*20*). The enrichment of crRNAs targeting host degradation genes in this region likely helps facilitate survival of the host, albeit to a lesser extent than targeting A1 and A2.

We additionally identified 36 Semi-Fit genes in T5, 22 of which had clear annotations: T5.014 (host genome nicking endonuclease), T5.037 (thioredoxin), rnh (Rnase H), nrdB (ribonucleoside diphosphate reductase small subunit), T5.102 (SIR2 deacetylase), T5.108 (replication origin binding protein), hegI (homing endonuclease), ligA (DNA ligase), D10 (replicative DNA helicase), D12 (palindromic endonuclease), D14 (Holliday junction resolvase), D15 (DNA exonuclease), D17 (straight tail fiber), D16 (tail fiber J), T5.139 (distal tail protein), D18-19 (tail length tape measure protein), T5.143 (tail assembly chaperone), N4 (major tail protein), T5.146 (tail terminator protein), T5.155 (terminase large subunit), sciB/T5.156 (terminase small subunit). An additional 3 Semi-Fit genes had ambiguous annotations: T5.035 (Ser-Thr Phosphatase), T5.147 (tail completion protein), T5.154 (endonuclease).

It should be noted that outside of the pre-early region in T5, we noticed a decrease in CRISPRi-ART consistency between different crRNAs targeting the same genes, especially at high MOIs. This means that many clearly-essential genes in T5 are omitted from Fit classification. For instance, in the core region of T5 between D2 and D20-21, many likely essential genes fall in the Semi-Fit category due to crRNA to crRNA variability (D18-D19 tape measure protein is a clear example). However inclusion of Semi-Fit genes introduces weaker-than-normal precision in analysis of the T5 genome too. For instance, leading to the likely faulty inclusion of T5.056 (*31*). Relative to other phages in this study, direct interpretation of Semi-Fit genes should be approached with extra caution during analysis of the T5 genome due to this decreased precision.

Outside of the pre-early region, we observed a clear enrichment of crRNAs targeting replication and virion proteins in the region between *D2* and *oad* (region from 63,000-111,000). In particular, of the 18 replication genes annotated as replicative in T5 that were not called Fit or Semi-Fit were *D2*, *ligB*, *pri*, and *D13*. In addition, only 6 of 16 virion proteins were not Fit or Semi-Fit. Of these, *T5.151* (hoc-like protein, known dispensable), *T5.036* (an Ambiguous annotation), *ltf* (L-tail fiber), and T5.136 (L-tail fiber attachment) are likely truly non-essential for infection of O-antigen deficient *E.coli* (*19*, *32*, *33*).

Finally, we observed 5 proteins of completely unknown function or roles in T5 with Fit status: T5.047. T5.084, T5.089, T5.090, and T5.114. Some of these proteins are among the most fit in our screens in T5 (T5.047 and T5.114). T5.114 colocalizes with another Fit gene, *D3*, suggestive of a confident, but unknown role of this two-gene cassette in the context of T5 infection. The gene *T5.047* is localized nearby, but not within, a region of known dispensability in T5 (*31*). These proteins might be a reflection of to-be-discovered mechanisms of how T5 interacts with *E.coli* DH10b or missing, non-structural components of the T5 lifecycle.

Supplementary Text S4. *Summary of Genome-wide screens in SUSP1.*

Our screens identified 20 Fit genes in non-model phage SUSP1: a*gp2*, a*gp6*, a*gp7*, *agp10*, *agp12*, *agp13*, *agp14*, *agp16*, *agp24*, *agp27*, *agp38*, *agp46*, *agp48*, *agp54*, *agp55*, *agp61*, *agp89*, *agp130*, *agp137*, and a*gp138*. As a non-model phage, SUSP1 has relatively little known about its biology beyond being a "superspreader phage" (*34*). In addition, SUSP1 displays significant homology to *Salmonella* phage FelixO1, a *Salmonella*-specific, but broad host range phage within species, recently identified to have complex genetic interactions with its host (*22*, *35*, *36*). Of the 138 coding annotations for SUSP1, 114 ultimately derive from FelixO1 via Mauve whole genome alignment and may reflect on this classical phage's biology (*21*, *22*).

Of the 19 Fit genes, 9 had clear- and 2 had ambiguous- annotations: a*gp2* (portal protein), a*gp6* (tail protease), a*gp7* (major capsid protein), a*gp12* (tail sheath), a*gp14* (tail assembly chaperone), a*gp16* (tape measure protein), a*gp46* (DNA polymerase), a*gp55* (exonuclease), a*gp61* (ribonucleoside-diphosphate reductase large subunit (NrdA)), a*gp13* (virion protein), a*gp48* (tail protein). These proteins primarily reflect structural features of the phage, but also include nucleotide metabolism (a*gp61*/*nrdA*) and replicative functions (a*gp46* and a*gp55*). The remaining 9 Fit genes are of entirely hypothetical function.

We additionally identified 12 genes of Semi-Fitness in at least one MOI: a*gp1* (terminase large subunit), a*gp15* (tail assembly chaperone), a*gp20* (baseplate spike protein), a*gp21* (baseplate wedge subunit), a*gp22* (baseplate wedge subunit), a*gp23* (baseplate protein), a*gp29* (thymidylate synthase), a*gp30* (dihydrofolate reductase), a*gp63* (ribonucleoside diphosphate reductase small subunit (NrdB)), a*gp17* (virion protein), a*gp18* (virion protein), and a*gp100* (hypothetical protein). 9 of these had clear annotations, 2 with unclear annotations and 1 with no known function.

Between the 31 Fit or Semi-Fit genes a very high percentage of the intuitively-essential SUSP1 genome was captured. Of the 19 predicted structural components (17 of them directly measured confirmed present in FelixO1 virions (*22*)), 10 appear Fit for SUSP1 and only miss proteins with ambiguous (*agp003*, *agp004*, *agp008*, *agp009*, *agp111*, *agp125*) or adsorptive functions that might have functional redundancy and thus dispensable against *E.coli* DH10B (*agp020*, *agp025*, *agp026*). 3 of 5 of the identifiable chaperonins appeared Fit as well as 4 of 6 identifiable replication genes (missing *agp040* encoding DNA Ligase and *agp051* encoding DNA primase). While SUSP1 is a non-model phage, our results corroborate that the high-level understanding of the important structural and replicative components of SUSP1 (and by extension, FelixO1) is well-understood.

Notably absent from our Fit or Semi-Fit genes are any genes related to lysis by SUSP1. Strangely, this is a consistent theme during the study of lysis in *Ounaviridae*, which have unique phenotypes surrounding lysis. SUSP1 (and FelixO1) release plasmids into the environment upon lysis of the infected cell. Model *Ounavirus*, FelixO1 (nonetheless SUSP1) to date lacks an identifiable holin (*22*), the primary essential component for cellular lysis by double-stranded DNA phages (*37*). The lysis aspect of the *Ounavirus* lifecycle remains a mystery.

Across the four phages assessed in this study, SUSP1 was the only phage to show strong phenotypes when targeting genes responsible for *de novo* nucleotide biosynthesis (*agp029*, *agp030*, *agp061*, and *agp063*) (*38*). Potentially, this too is related to the "superspreader" phenotypes documented with SUSP1 (*34*). Because SUSP1 doesn't digest the host genome or plasmids during the course of infection, SUSP1 might be comparatively reliant on *de novo* nucleotide biosynthesis during replication. Although these genes have homologs in phage T4 (*frd*, *td*, *nrdA*, and *nrdB*), these genes do not show up as fit in T4. Potentially, one of the trends in the evolution of phages with larger genomes is the ability to acquire "building blocks" via multiple strategies.

Finally, our collective results identify 10 genes in SUSP1 of completely hypothetical function displaying Fit or Semi-Fit phenotypes: *agp10*, *agp24*, *agp27*, *agp38*, *agp54*, *agp89*, *agp100*, *agp130*, *agp137*, and *agp138*. On average, these genes were smaller on average and were dispersed across the SUSP1 genome. Occasionally, as the case for *agp130*, these genes were among the top scoring in our SUSP1 Fit screens.

To identify candidates that may reflect essential host-adaptation to *E.coli* relative to *Salmonella* we looked for genes that were Fit in SUSP1, but were not in the FelixO1 genome. We identified 3 such genes: *agp038*, *agp137*, and *agp138*. One particularity is the location of *agp137* and *agp138*, which occur amidst the tRNA-encoding region of SUSP1. Potentially these genes in the auxiliary genome of *Ounaviridae* reflect an adaptation to host defenses that target tRNAs.

Supplementary Text S5. *Summary of Genome-wide screens in PTXU04.*

Our screens identified 17 Fit genes in singleton phage PTXU04: *gp1*, *gp7*, *gp8*, *gp9*, *gp10*, *gp11*, *gp19*, *gp20*, *gp22*, *gp23*, *gp24*, *gp25*, *gp28*, *gp38*, *gp40*, *gp41*, and *gp92*. PTXU04 was originally reported as a "singleton phage" and we find it to have only marginal similarity to other isolated phages: in our network graph, PTXU04 only shares homology of one of its proteins with neighboring phages (Fig. 4A) (*4*). Because of this distant relation to other studied phages, annotation of PTXU04 is comparatively poor and is predominantly encoded by genes of unknown function. All functional annotations generally need to be approached with skepticism.

Of the 17 fit genes identified in this study, only 5 had confidently inferable function with hhpred: *gp1*, *gp7*, *gp10*, *gp38*, and *gp41*, encoding small terminase subunit, portal protein, putative major capsid protein (*mcp*), single-stranded DNA binding protein (SSB), and replicative DNA helicase, respectively. An additional 4 genes had ambiguous but weak function: *gp9*, *gp11*, *gp28*, and *gp92*, encoding putative head scaffolding protein, virion protein, RNA binding protein, and endonuclease respectively. Another gene, *gp25*, harbored a "Large Polyvalent Domain 38" (LPD38) annotation at its C-terminus (*39*). While its predicted size and intrinsic disorder suggest that it's a polyprotein, no other domains of confidence were inferrable in this large protein. Polyvalent proteins play an intriguing role in phage biology and are predicted to play enigmatic roles in the phage-host arms race (*39*).

We additionally identified 13 genes of Semi-Fitness in at least one MOI: *gp3*, *gp16*, *gp17*, *gp18*, *gp22*, *gp28*, *gp34*, *gp35*, *gp37*, *gp42*, *gp44*, *gp58*, *gp60*, *gp74*, and *gp89*. Only 2 had confidently inferable functions with hhpred (*13*): *gp35* and *gp44*, encoding DNA polymerase and DNA primase, respectively. Another 4 genes encoded inferred structural components, but lacked confident annotations: *gp3*, *gp16*, *gp17*, *gp18*.

In aggregate, these screens reveal a reliable trend with the PTXU04 genome. Consistent with the model phages, ostensible essential genes in PTXU04 appear to colocate in several regions of the genome: *gp1*- *gp11*, gp16-*gp25, and gp34-gp44*. However, we observed notable omissions of many of the expected essential genes encoded within. For instance, *gp2*, *gp4*, and *gp30* encode likely essential genes encoding large terminase, tail fiber, and lysozyme encoded within otherwise Fit regions that display subthreshold fitness. Some of these genes might be dispensable in this infection context (ex. Tail fiber). Others, such as *gp30* might merely exhibit "true" crRNA to crRNA variability, as its maximal fitness guide performs better, limiting their fitness in this context.

Finally, maybe the most curious outcome of these screens is the clear existence of a highly Fit series of genes from *gp16*-*gp25*, the vast majority of which have no identifiable function. This result suggests a probable deviation for PTXU04 from other phages in terms of its lifecycle and might provide a clear path for characterization of the enigmatic LPD domains of bacterial viruses.

**Supplementary Figures:**

**Fig. S1. Guide RNA Design for CRISPRi-ART RBS Tiling.**

**Fig. S2. Oligo pool design.**

**Fig. S3. Comparison of CRISPRi-ART to other measurements of essentiality.**

**Fig. S4. Antisense vs sense crRNA fitnesses in *E. coli* RBS-targeting library.**

**Fig. S5. Quantitative Western of endogenous nonessential operon and qPCR measurements of targeted transcripts.**

**Fig. S6. T4 CRISPRi-ART Plaque Assays.**

**Fig. S7. T4 dCas12a CRISPRi Plaque Assays.**

**Fig. S8. T4 dCas9 CRISPRi Plaque Assays.**

**Fig. S9. EdH4 CRISPRi-ART Plaque Assays.**

**Fig. S10. Goslar CRISPRi-ART Plaque Assays.**

**Fig. S11. Lambda CRISPRi-ART Plaque Assays.**

**Fig. S12. M13 CRISPRi-ART Plaque Assays.**

**Fig. S13 MM02 CRISPRi-ART Plaque Assays.**

**Fig. S14. MS2 CRISPRi-ART Plaque Assays.**

**Fig. S15. N4 CRISPRi-ART Plaque Assays.**

**Fig. S16. PTXU04 CRISPRi-ART Plaque Assays.**

**Fig. S17. SUSP1 CRISPRi-ART Plaque Assays.**

**Fig. S18. T5 CRISPRi-ART Plaque Assays.**

**Fig. S19. T7 CRISPRi-ART Plaque Assays.**

**Fig. S20. Genome-wide CRISPRi-ART Fitness for Phage T4.**

**Fig. S21. Genome-wide CRISPRi-ART Fitness for Phage T5.**

**Fig. S22. Genome-wide CRISPRi-ART Fitness for Phage SUSP1.**

**Fig. S23. Genome-wide CRISPRi-ART Fitness for Phage PTXU04.**

**Fig. S24. crRNA Fitness Distributions for Top 10 Fit T4 Genes.**

**Fig. S25. crRNA Fitness Distributions for Top 10 Fit Phage T5 Genes.**

**Fig. S26. crRNA Fitness Distributions for Top 10 Fit Phage SUSP1 Genes.**

**Fig. S27. crRNA Fitness Distributions for Top 10 Fit Phage PTXU04 Genes.**

**Fig. S28. crRNA fitness versus plaque size for T5 *D20-21* (*mcp*).**

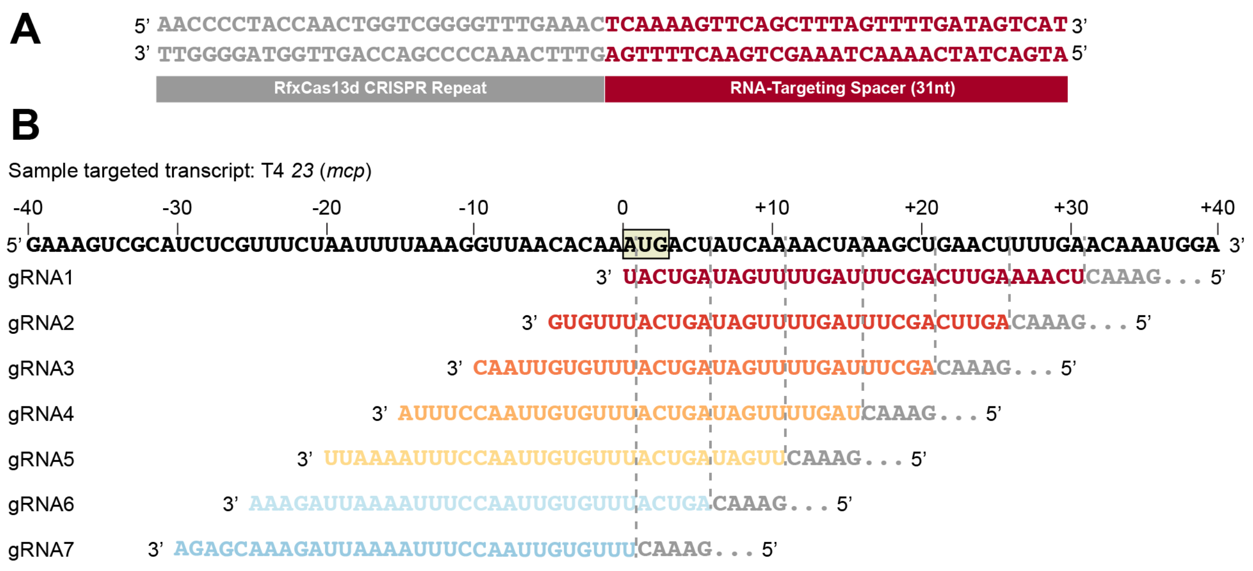


**Fig. S1. Guide RNA Design for CRISPRi-ART RBS Tiling.**

All examples in fig. S1 refer to crRNAs (referred to here as guide RNAs, or gRNAs) that target Phage T4 *23* (*mcp*) (**A**). Example T4 *mcp* targeting gRNA1 encoded in dRfxCas13d plasmid consisting of a CRISPR repeat (gray) and a 31 nucleotide spacer (red, reverse complement of targeted transcript) (**B**). For RBS-tiling designs, gRNAs are designed using 31 nucleotide spacers targeting every 5 nucleotides along a transcript from +31 to +1 starting positions for gRNA1 to gRNA7 respectively. Mature RNA targeting sequences are highlighted in rainbow. Partial CRISPR repeats are shown in gray. Targeted transcript is shown in black with the start codon highlighted.

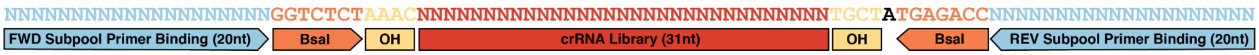


**Fig. S2. Oligo pool design for crRNA libraries.**

All oligos in the crRNA oligo pools encode two 20 nt subpool-specific priming sites to facilitate selective amplification of distinct subpools, two BsaI sites with appropriate overhangs to clone into oligo library entry vectors, and 31 nt crRNAs.

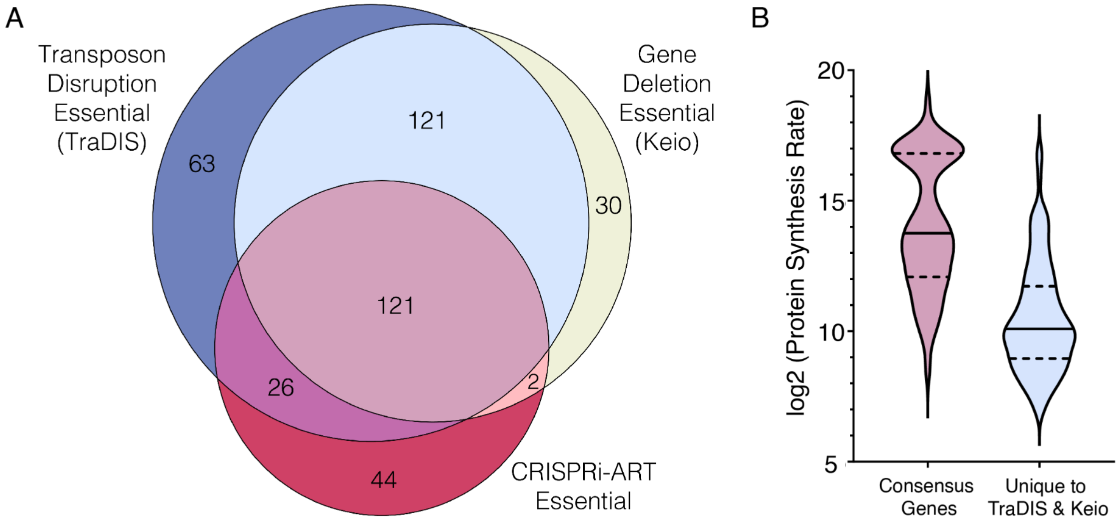


**Fig. S3. Comparison of CRISPRi-ART to other measurements of essentiality in context of protein synthesis rate.**

A. Venn diagram comparing essential genes identified by CRISPRi-ART, KEIO, and TraDIS. The diagram illustrates the significant intersection of genes with negative fitness impacts across three techniques: CRISPRi-ART, where genes were deemed essential if at least 3 RBS-targeting gRNAs resulted in an average $\log_2$FC of <-2; KEIO-based homologous recombination; and TraDIS-based transposon sequencing. B. Violin plot displaying the log2 protein synthesis rates for genes. Pink represents genes effectively targeted by all three techniques (Consensus Genes, n=121), while light blue denotes genes where CRISPRi-ART crRNAs did not yield a fitness defect meeting the criteria to be deemed “significantly impacted” (Unique to TraDIS and Keio, n=121). The width of each violin indicates the density of the data at different synthesis rates, with wider sections representing higher densities of data points. Genes called by all three techniques have an average protein synthesis rate (in units of molecules produced per generation) of (14.0) while genes that are missed by our technique have an average synthesis rate of (10.4). SD is represented by dashed lines in violin plots.

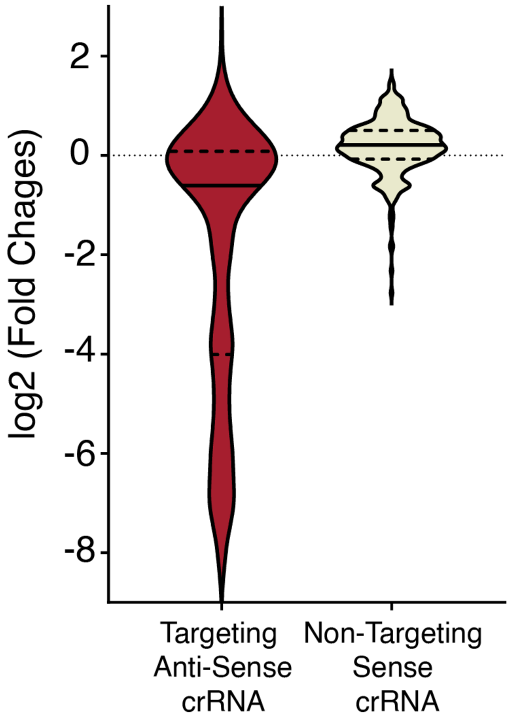


**Fig. S4. Antisense vs sense crRNA fitnesses in *E. coli* RBS-targeting library.**

Violin plot displaying the $log_2FC$ for crRNA targeting essential genes in the *E. coli* genome. Red represents antisense crRNAs targeting the RBS region of a gene of interest (Targeting Anti-Sense crRNA) n=997, while tan denotes genes non-targeting sense crRNA, targeting the center of the sense strand of the RBS region (Non-Targeting Sense crRNA) n=325. The width of each violin indicates the density of the data at different synthesis rates, with wider sections representing higher densities of data points. crRNA targeting the RBS of essential genes have an average $log_2FC$ of (-1.95) while crRNA targeting the sense strand of the RBS region have an average $log_2FC$ of (0.12). SD is represented by dashed lines in violin plots.

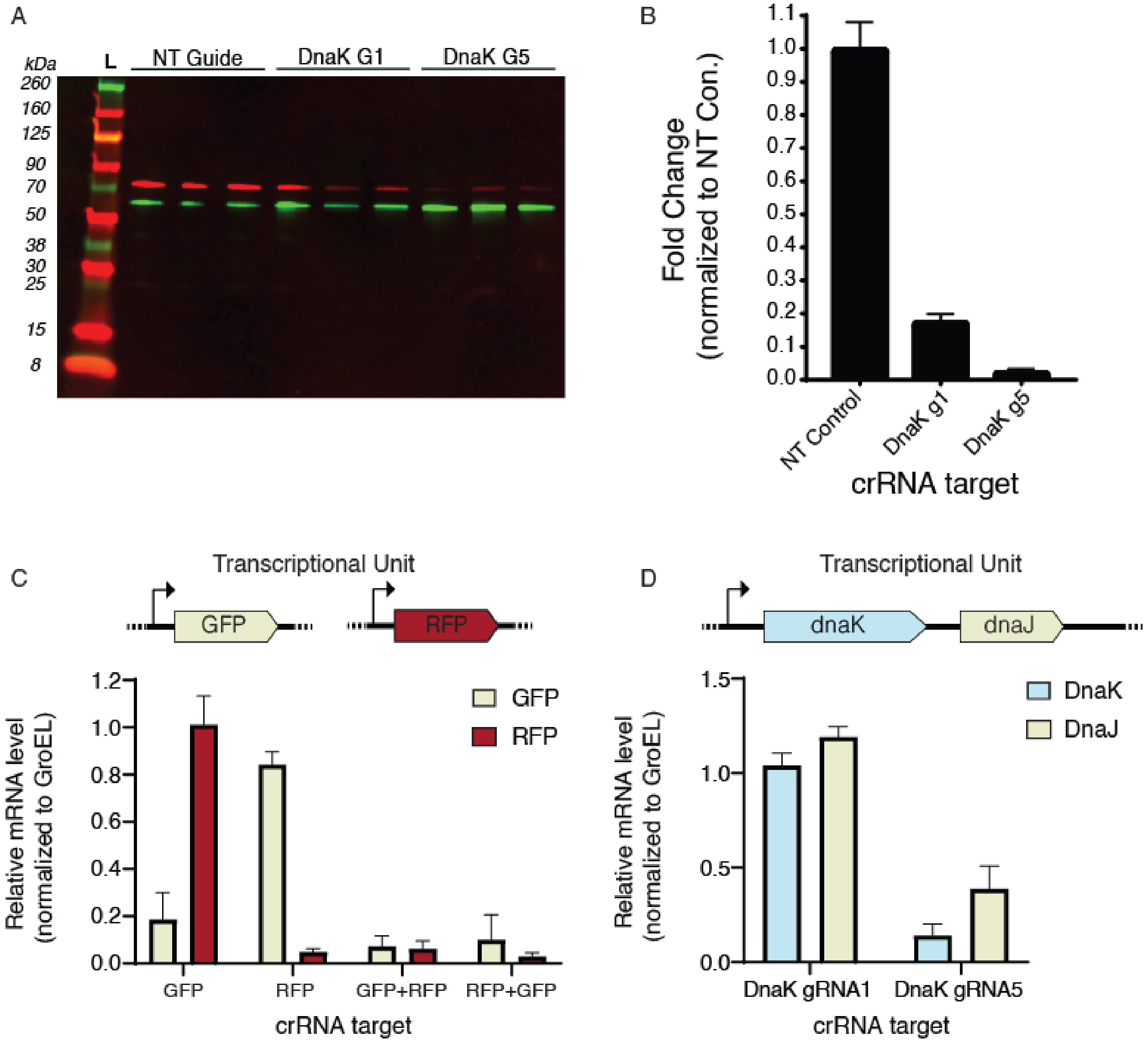
A
kDa
L
NT Guide
DnaK G1
DnaK G5
260
160
125
90
70
50
38
30
25
15
8
B
Fold Change
(normalized to NT Con.)
1.1
1.0
0.9
0.8
0.7
0.6
0.5
0.4
0.3
0.2
0.1
0.0
NT Control
DnaK g1
DnaK g5
crRNA target
C
Transcriptional Unit
GFP
RFP
Relative mRNA level
(normalized to GroEL)
1.2
1.0
0.8
0.6
0.4
0.2
0.0
GFP
RFP
GFP
RFP
GFP+RFP
RFP+GFP
crRNA target
D
Transcriptional Unit
dnaK
dnaJ
Relative mRNA level
(normalized to GroEL)
1.5
1.0
0.5
0.0
DnaK
DnaJ
DnaK gRNA1
DnaK gRNA5
crRNA target

**Fig. S5. Quantitative Western of endogenous nonessential operon and qPCR measurements of targeted transcripts.**

A. Western blot showing DnaK protein quantity in triplicate as targeted by three gRNA (NT,gRNA1,gRNA5) shown in red. GroEL (E.Coli Protein chaperone GroEL) shown as loading control (green) B. Densitometric analysis of the bands from (A) normalized to GroEL. The relative protein levels of DnaK are presented as fold change compared to NT guide control. Data represent mean ± SD of three biological replicates. C.D.Relative transcript abundance (normalized to non-targeting crRNA) of the indicated targets in *E. coli* cells after cells reach an $OD_{600}$ ~0.5 of dCas13d plasmid with corresponding crRNA, measured by RT-qPCR. Error bars indicate mean ± s.d. of three biological replicates.

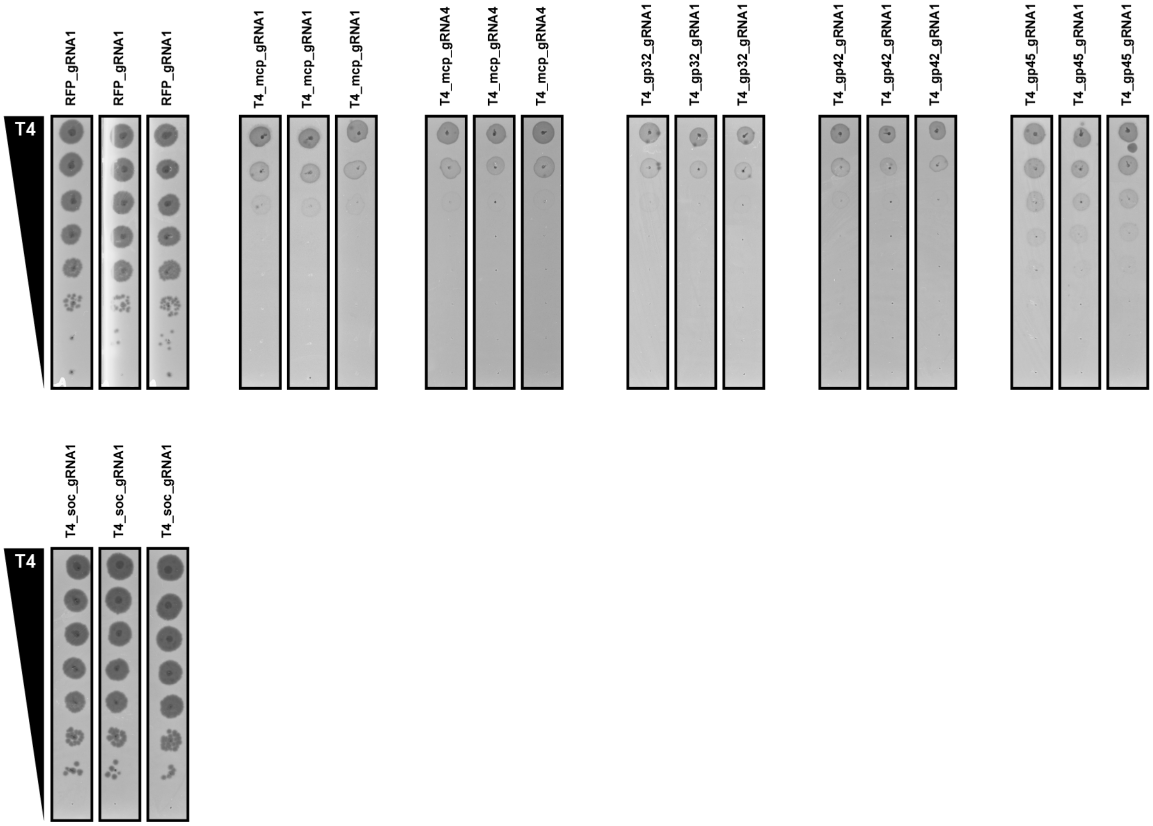


**Fig. S6. T4 CRISPRi-ART Plaque Assays.**

Plaque assays for CRISPRi-ART-mediated phage defense when targeting phage T4 RBS with dRfxCas13d. A guide targeting RFP is provided as a negative control. dRfxCas13d experiments were expressed using +20nM aTc. Data shown are for 3 biological replicates.

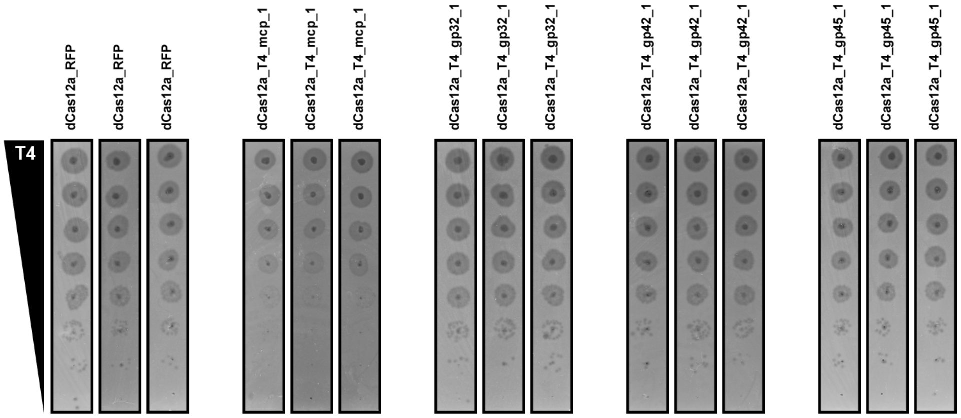


**Fig. S7. T4 dLbCas12a CRISPRi Plaque Assays.**

Plaque assays for CRISPRi-mediated phage defense when targeting phage T4 CDS with dRfxCas13d. A guide targeting RFP is provided as a negative control. dLbCas12a experiments were expressed using +10nM aTc. Data shown are for 3 biological replicates.

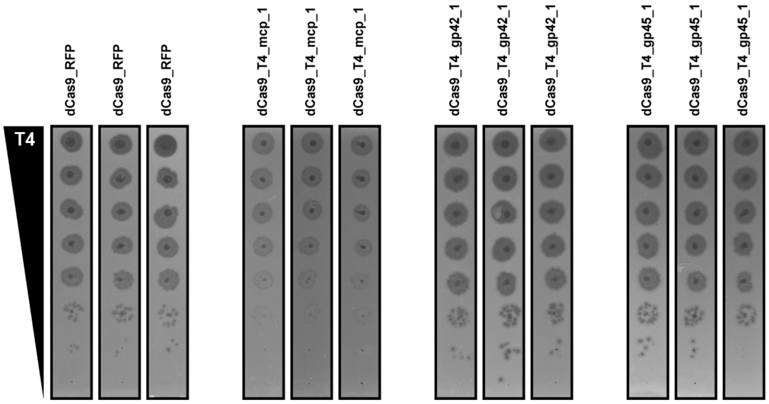


**Fig. S8. T4 dSpyCas9 CRISPRi Plaque Assays.**

Plaque assays for CRISPRi-mediated phage defense when targeting phage T4 CDS with dRfxCas13d. A guide targeting RFP is provided as a negative control. dSpyCas9 experiments were expressed using +20nM aTc. Data shown are for 3 biological replicates.

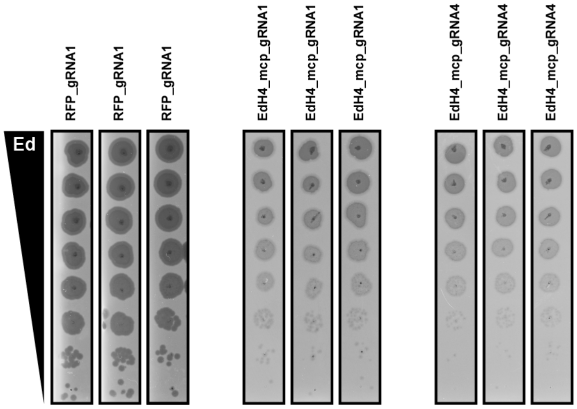


**Fig. S9. EdH4 CRISPRi-ART Plaque Assays.**

Plaque assays for CRISPRi-ART-mediated phage defense when targeting phage EdH4 RBS with dRfxCas13d. A guide targeting RFP is provided as a negative control. dRfxCas13d experiments were expressed using +100nM aTc. Data shown are for 3 biological replicates.

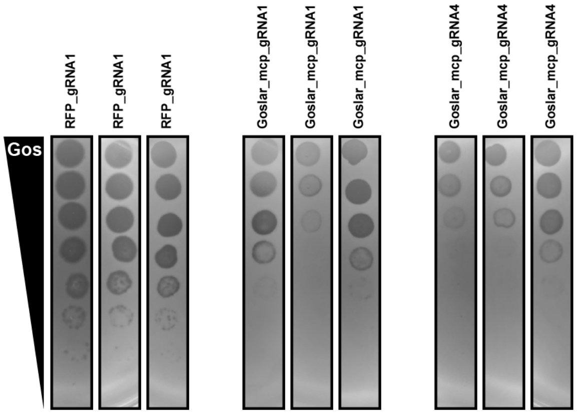


**Fig. S10. Goslar CRISPRi-ART Plaque Assays.**

Plaque assays for CRISPRi-ART-mediated phage defense when targeting phage Goslar RBS with dRfxCas13d. A guide targeting RFP is provided as a negative control. dRfxCas13d experiments were expressed using +100nM aTc. Data shown are for 3 biological replicates.

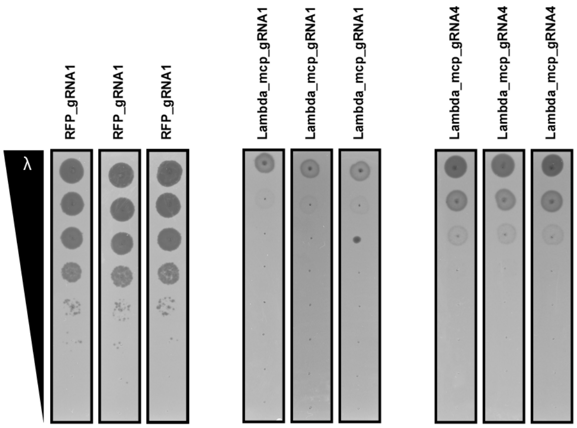


**Fig. S11. Lambda CRISPRi-ART Plaque Assays.**

Plaque assays for CRISPRi-ART-mediated phage defense when targeting phage Lambda RBS with dRfxCas13d. A guide targeting RFP is provided as a negative control. dRfxCas13d experiments were expressed using +20nM aTc. Data shown are for 3 biological replicates.

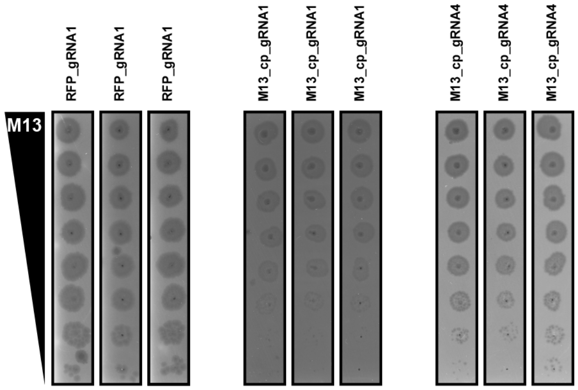


**Fig. S12. M13 CRISPRi-ART Plaque Assays.**

Plaque assays for CRISPRi-ART-mediated phage defense when targeting phage M13 RBS with dRfxCas13d. A guide targeting RFP is provided as a negative control. dRfxCas13d experiments were expressed using +100nM aTc and +1mM $CaCl_2$. Data shown are for 3 biological replicates.

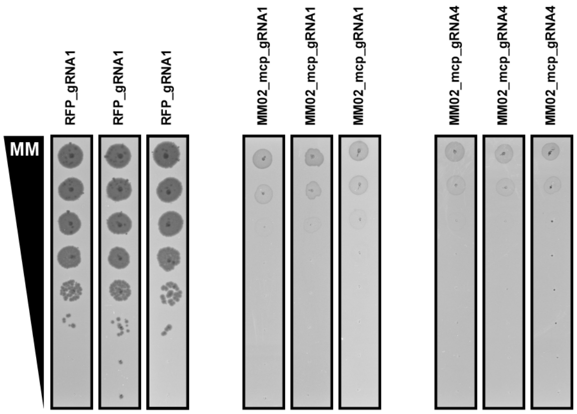


**Fig. S13 MM02 CRISPRi-ART Plaque Assays.**

Plaque assays for CRISPRi-ART-mediated phage defense when targeting phage MM02 RBS with dRfxCas13d. A guide targeting RFP is provided as a negative control. dRfxCas13d experiments were expressed using +20nM aTc. Data shown are for 3 biological replicates.

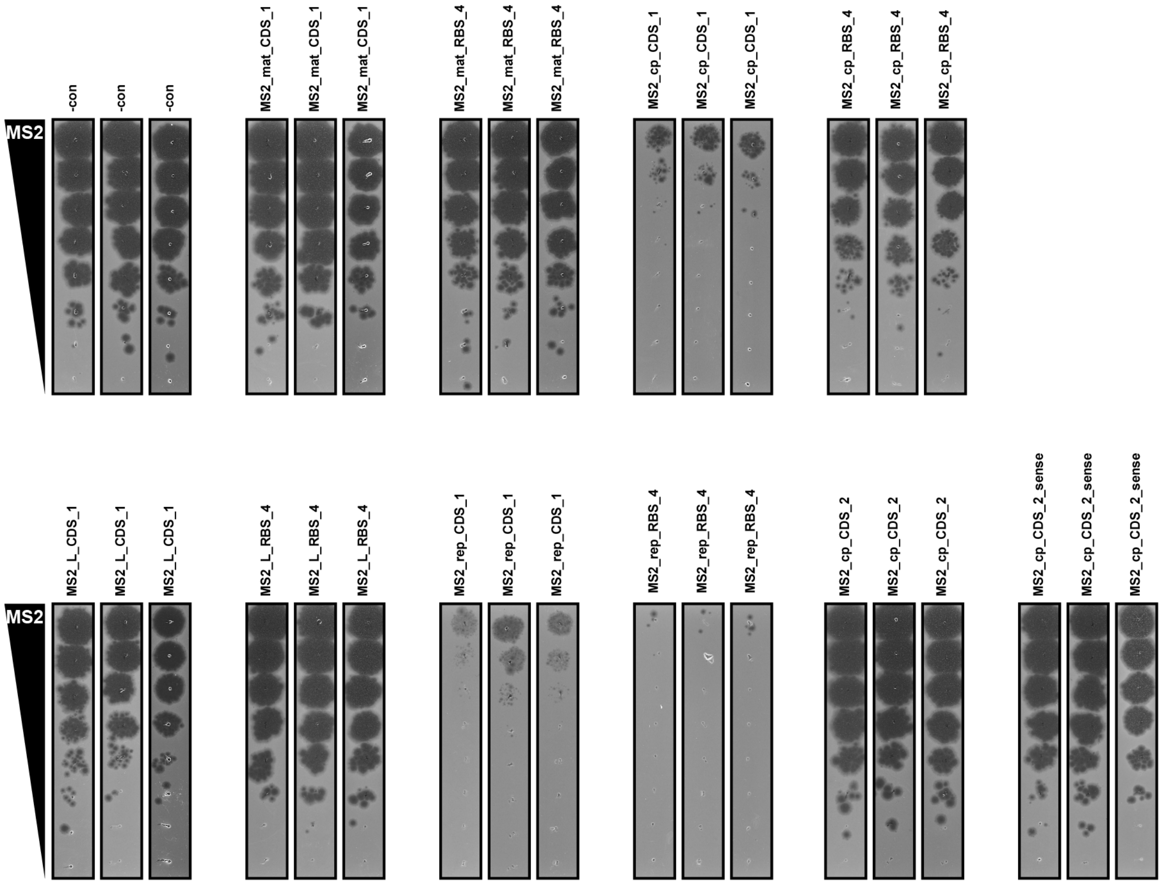


**Fig. S14. MS2 CRISPRi-ART Plaque Assays.**

Plaque assays for CRISPRi-ART-mediated phage defense when targeting phage MS2 RBS with dRfxCas13d. A guide targeting RFP is provided as a negative control. dRfxCas13d experiments were expressed using +100nM aTc and +1mM $CaCl_2$. Data shown are for 3 biological replicates.

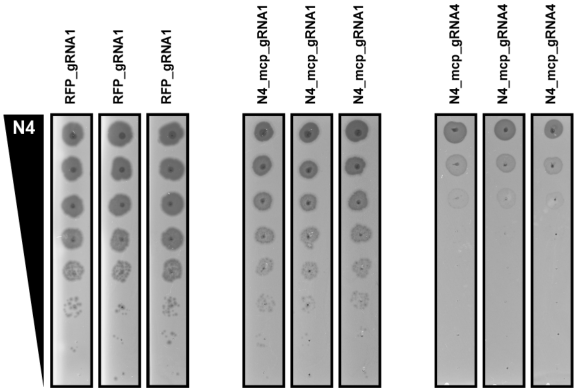


**Fig. S15. N4 CRISPRi-ART Plaque Assays.**

Plaque assays for CRISPRi-ART-mediated phage defense when targeting phage N4 RBS with dRfxCas13d. A guide targeting RFP is provided as a negative control. dRfxCas13d experiments were expressed using +100nM aTc. Data shown are for 3 biological replicates.

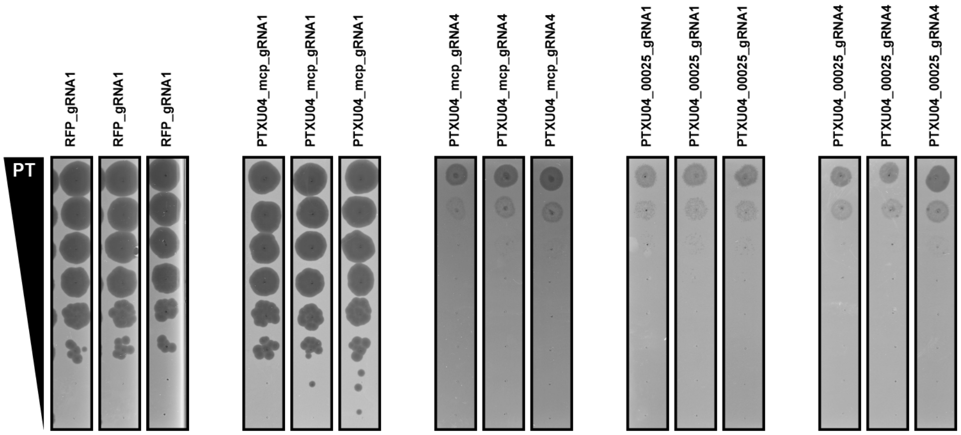


**Fig. S16. PTXU04 CRISPRi-ART Plaque Assays.**

Plaque assays for CRISPRi-ART-mediated phage defense when targeting phage PTXU04 RBS with dRfxCas13d. A guide targeting RFP is provided as a negative control. dRfxCas13d experiments were expressed using +100nM aTc. Data shown are for 3 biological replicates.

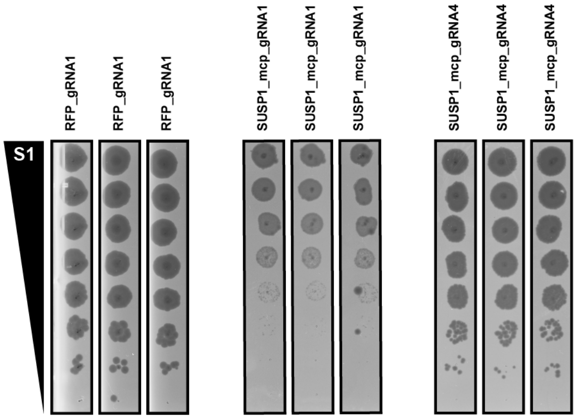


**Fig. S17. SUSP1 CRISPRi-ART Plaque Assays.**

Plaque assays for CRISPRi-ART-mediated phage defense when targeting phage SUSP1 RBS with dRfxCas13d. A guide targeting RFP is provided as a negative control. dRfxCas13d experiments were expressed using +100nM aTc. Data shown are for 3 biological replicates.

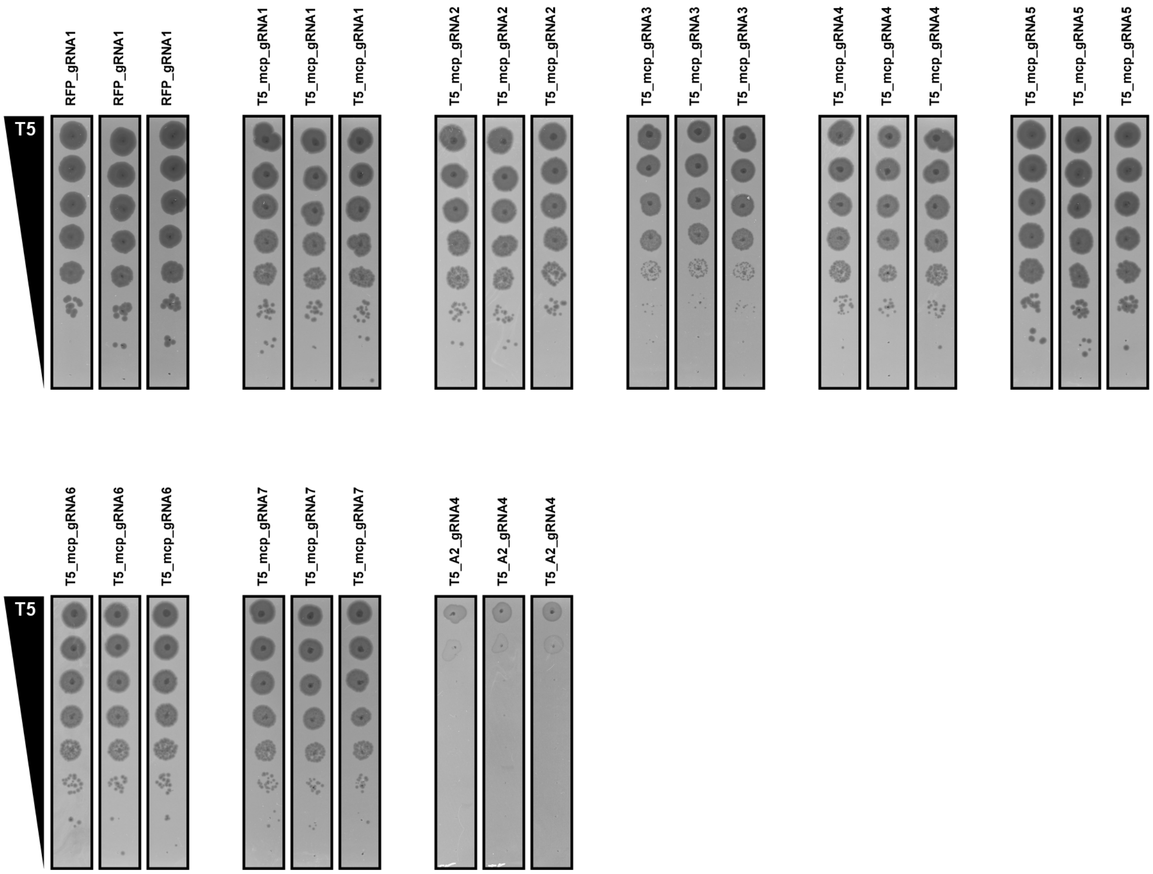


**Fig. S18. T5 CRISPRi-ART Plaque Assays.**

Plaque assays for CRISPRi-ART-mediated phage defense when targeting phage T5 RBS with dRfxCas13d. A guide targeting RFP is provided as a negative control. dRfxCas13d experiments were expressed using +100nM aTc. Data shown are for 3 biological replicates.

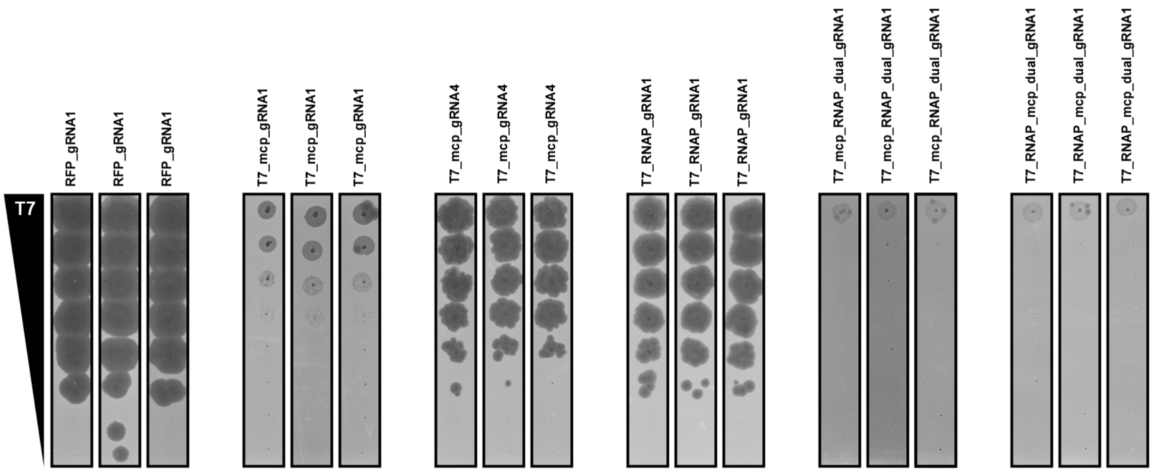


**Fig. S19. T7 CRISPRi-ART Plaque Assays.**

Plaque assays for CRISPRi-ART-mediated phage defense when targeting phage T7 RBS with dRfxCas13d. A guide targeting RFP is provided as a negative control. dRfxCas13d experiments were expressed using +100nM aTc. Data shown are for 3 biological replicates.

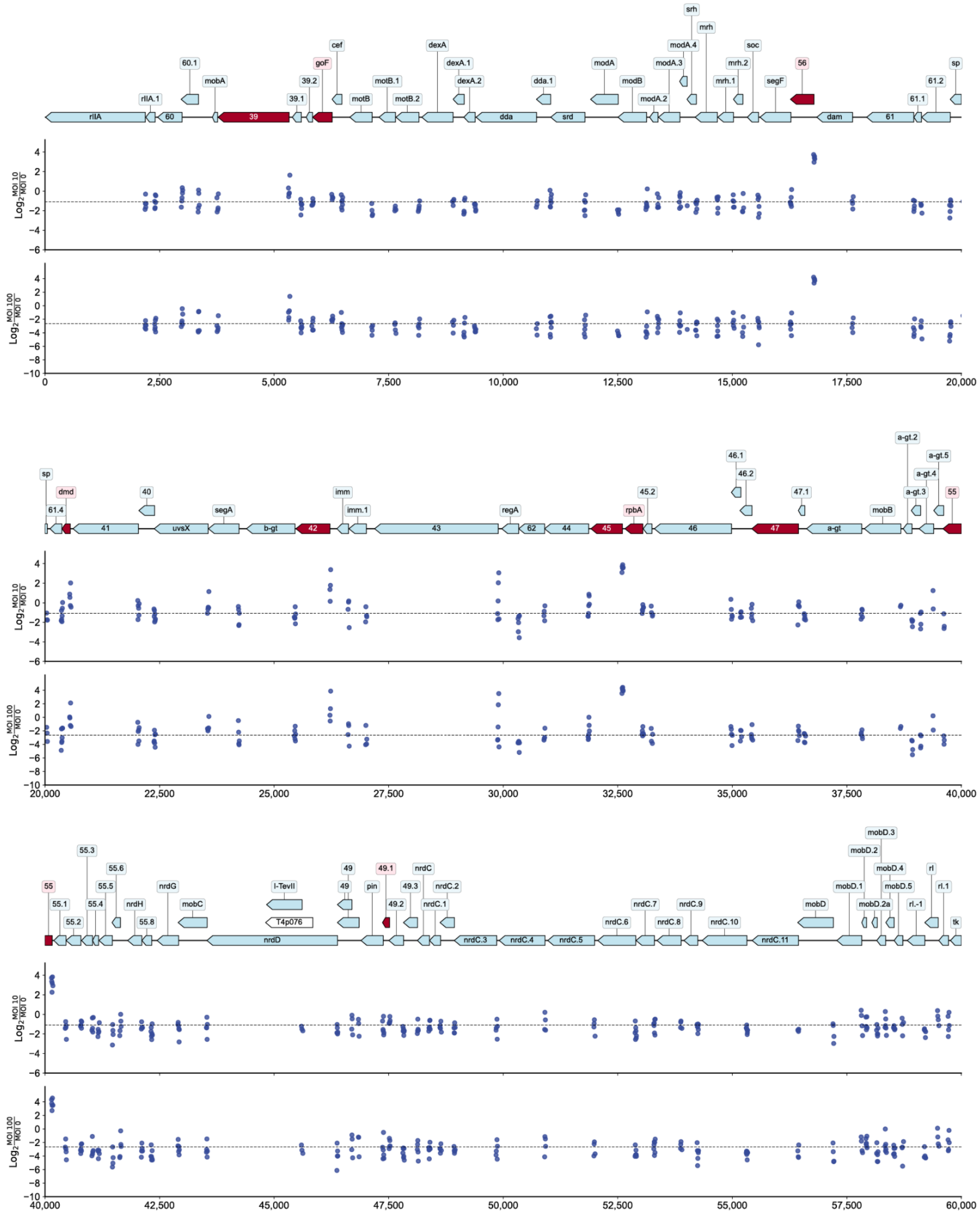


**Fig. S20. Genomewide ddCas13d-Fitness for Phage T4 (continued below)**

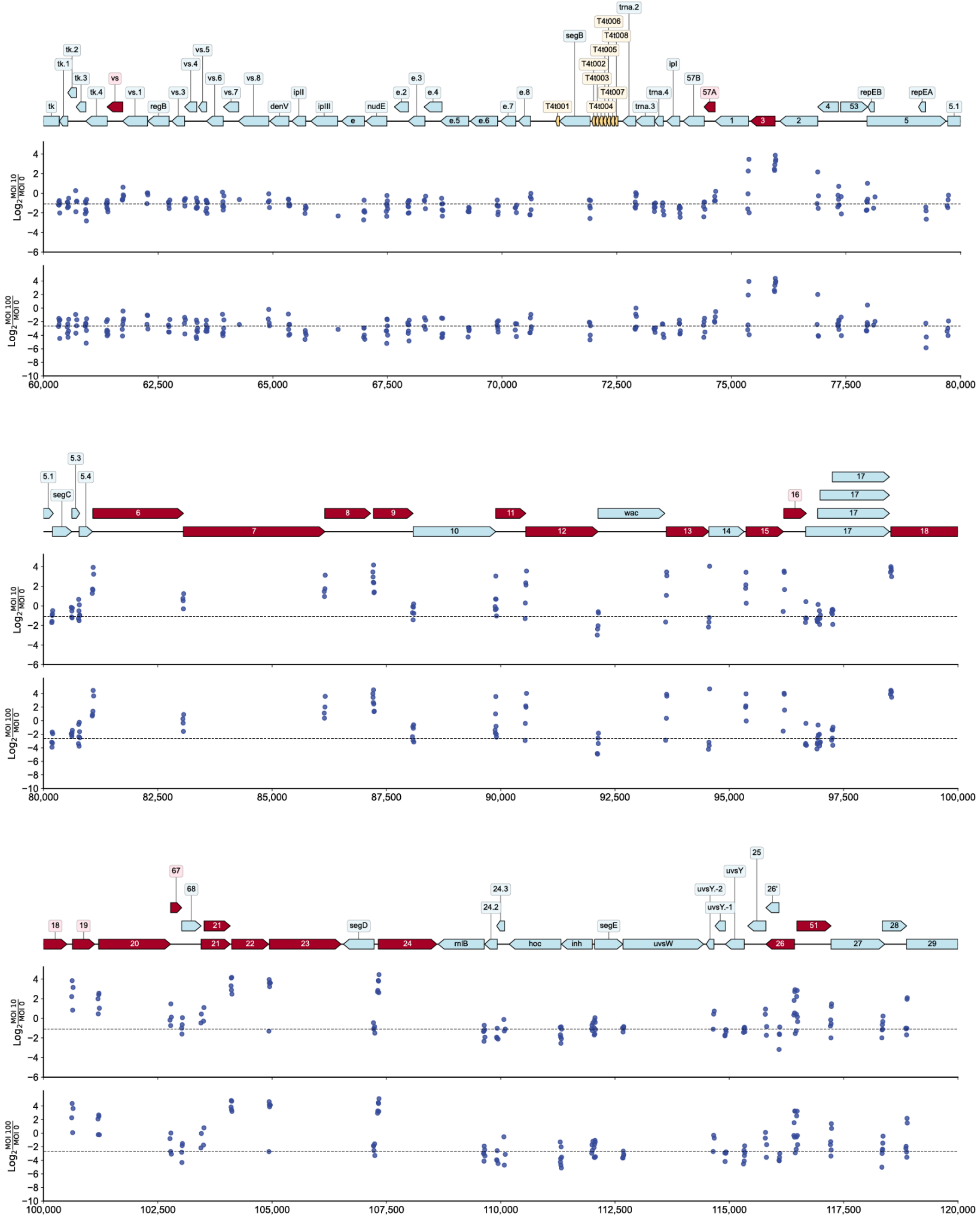


**Fig. S20. Genomewide ddCas13d-Fitness for Phage T4 (continued below)**

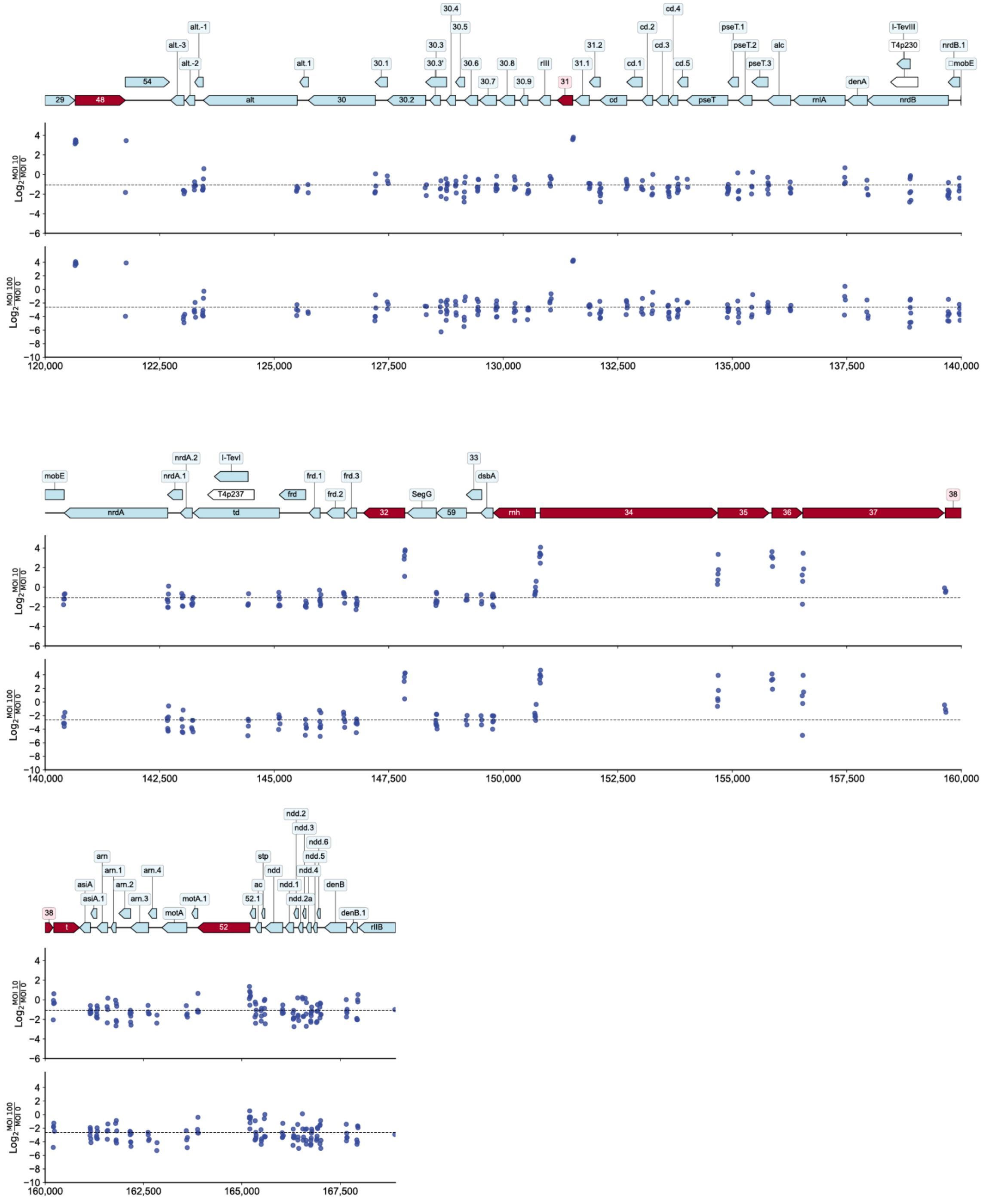


**Fig. S20. Genomewide ddCas13d-Fitness for Phage T4**

(Top track) Gene overview of phage T4 (NC_000866.4). Coding genes are shown in light blue and highlighted in red if they met significance thresholds (Methods). tRNAs are shown in orange. Other noncoding genetic elements are shown in white. (Middle track) ddCas13d gene knockdown fitness when targeting T4 genes during infection at

10 MOI. Median guide fitness is shown with a dashed line. (Bottom track) ddCas13d gene knockdown fitness when targeting T4 genes during infection at 100 MOI. Median guide fitness is shown with a dashed line.

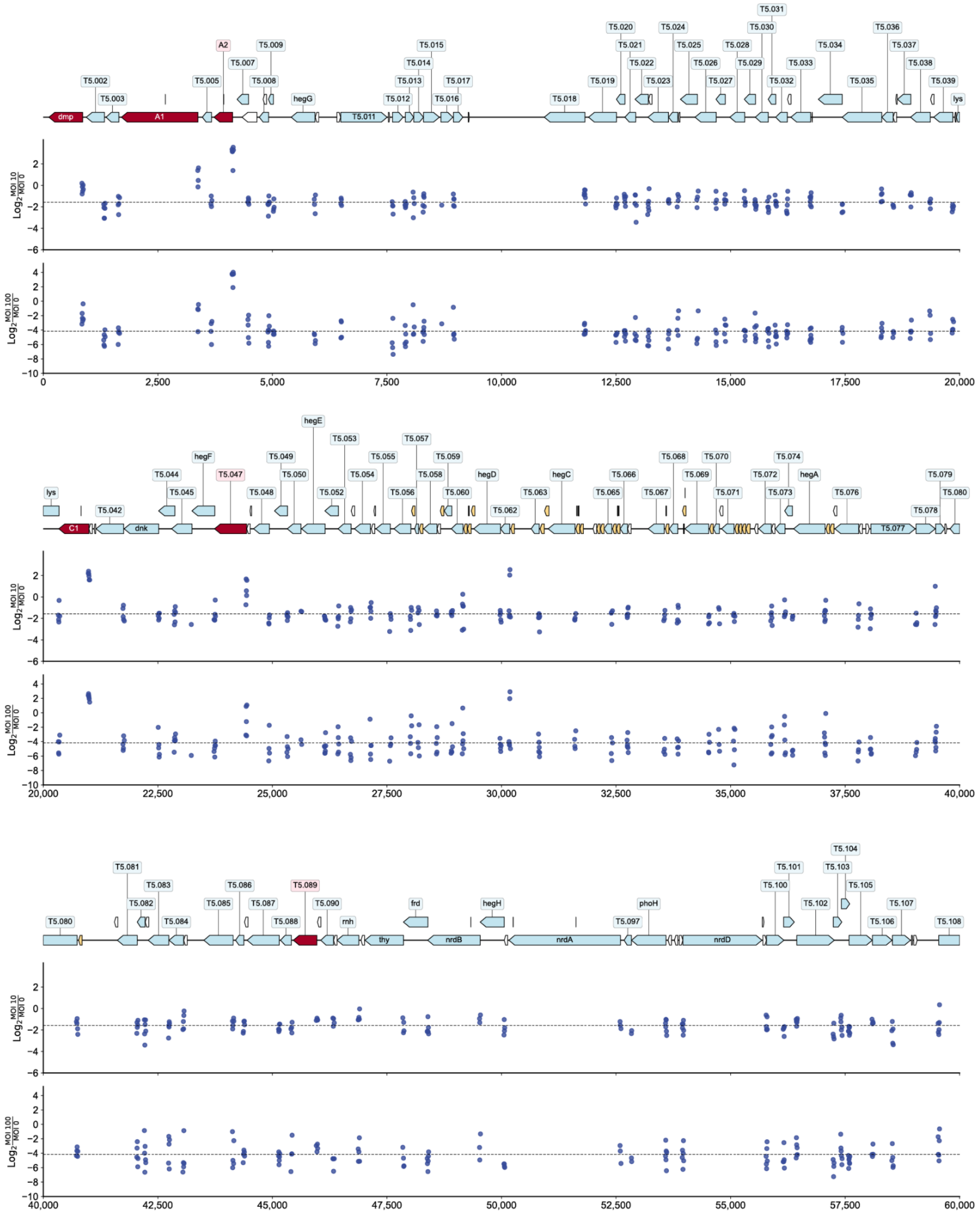


**Fig. S21. Genomewide ddCas13d-Fitness for Phage T5 (continued below)**

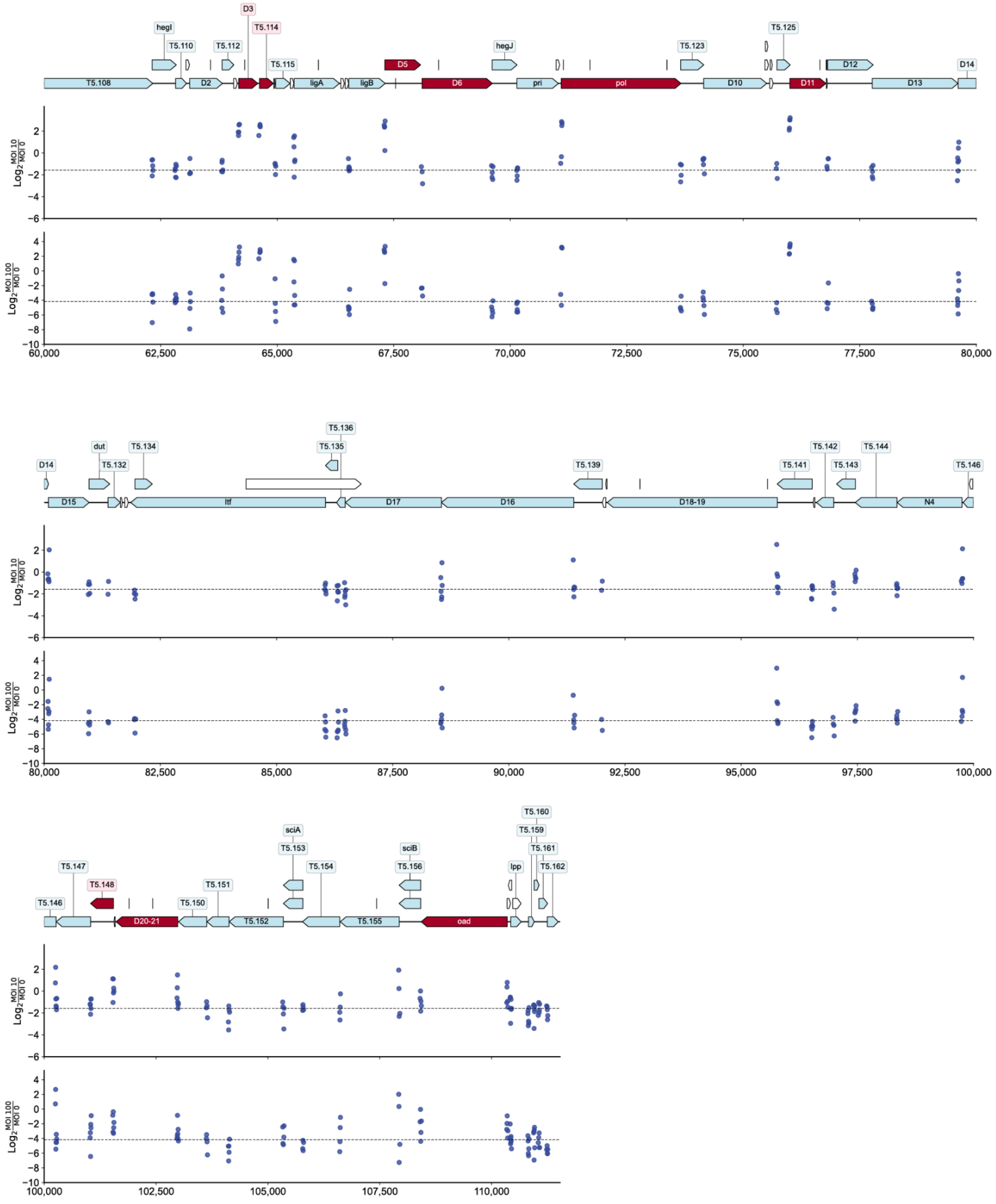


**Fig. S21. Genomewide ddCas13d-Fitness for Phage T5 (continued below)**

**Fig. S21. Genomewide ddCas13d-Fitness for Phage T5**

(Top track) Gene overview of phage T5 (NC_005859). Coding genes are shown in light blue and highlighted in red if they met significance thresholds (Methods). tRNAs are shown in orange. Other noncoding genetic elements are shown in white. (Middle track) ddCas13d gene knockdown fitness when targeting T5 genes during infection at 10 MOI. Median guide fitness is shown with a dashed line. (Bottom track) ddCas13d gene knockdown fitness when targeting T5 genes during infection at 100 MOI. Median guide fitness is shown with a dashed line.

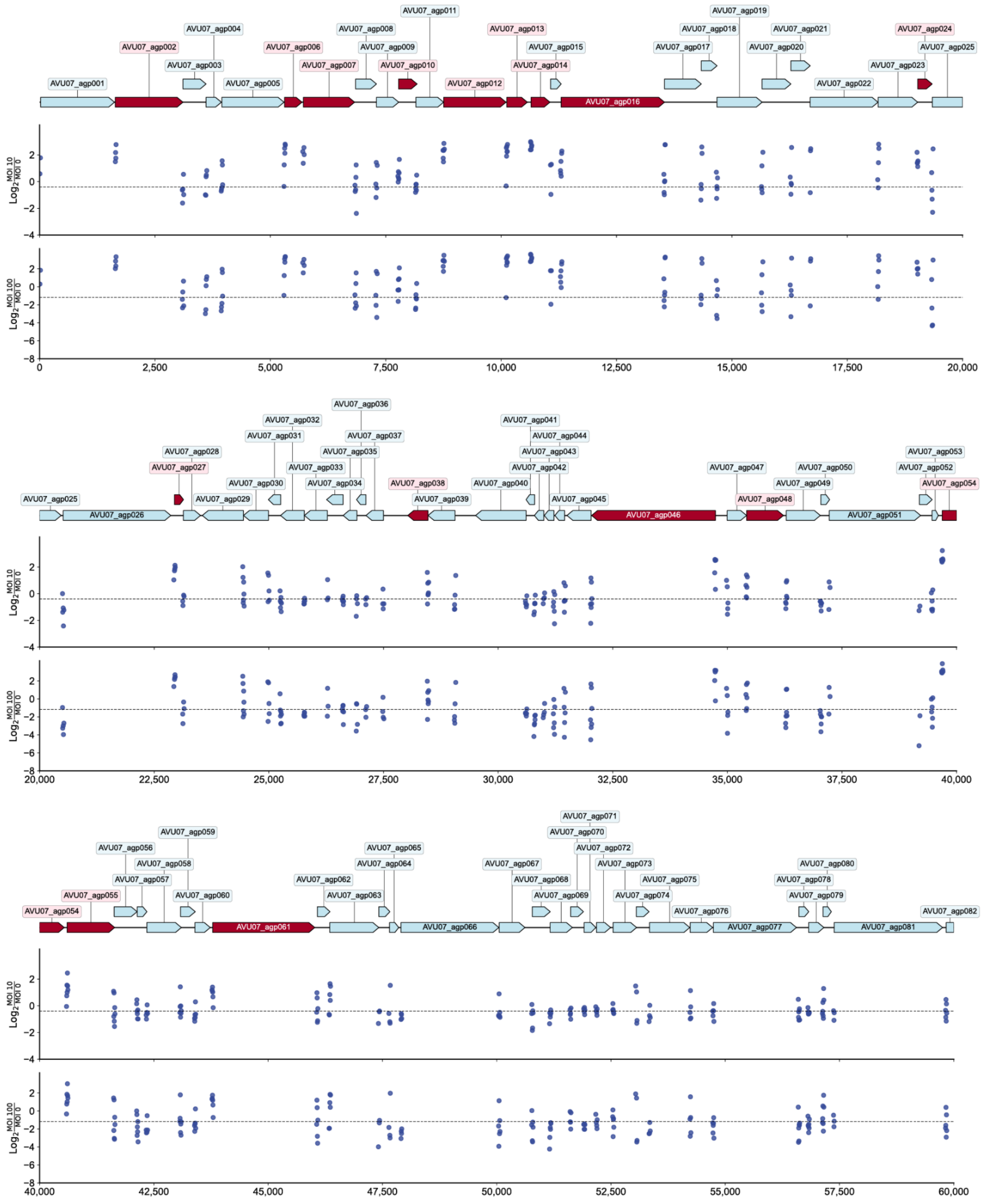


**Fig. S22. Genomewide ddCas13d-Fitness for Phage SUSP1 (continued below)**

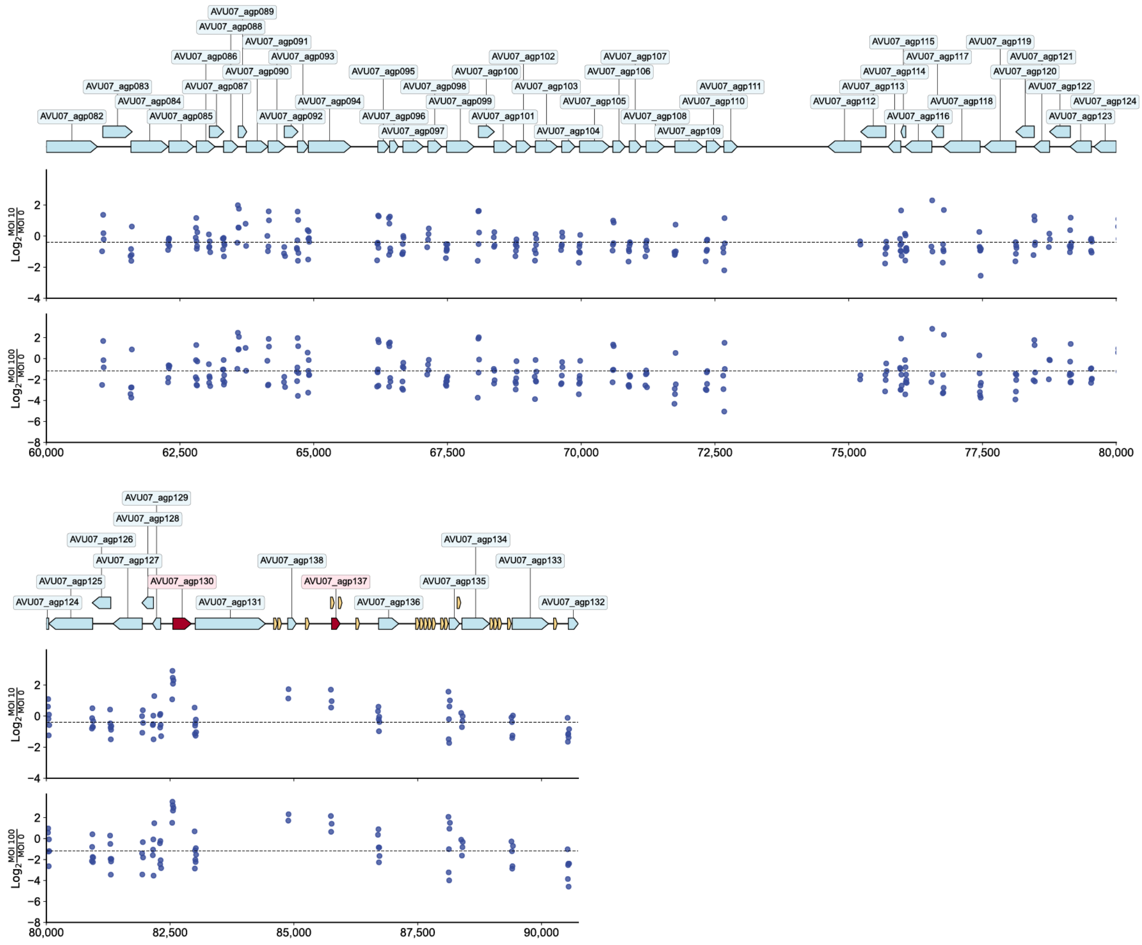


**Fig. S22. Genomewide ddCas13d-Fitness for Phage SUSP1**

(Top track) Gene overview of phage SUSP1 (NC_028808). Coding genes are shown in light blue and highlighted in red if they met significance thresholds (Methods). tRNAs are shown in orange. Other noncoding genetic elements are shown in white. (Middle track) ddCas13d gene knockdown fitness when targeting SUSP1 genes during infection at 10 MOI. Median guide fitness is shown with a dashed line. (Bottom track) ddCas13d gene knockdown fitness when targeting SUSP1 genes during infection at 100 MOI. Median guide fitness is shown with a dashed line.

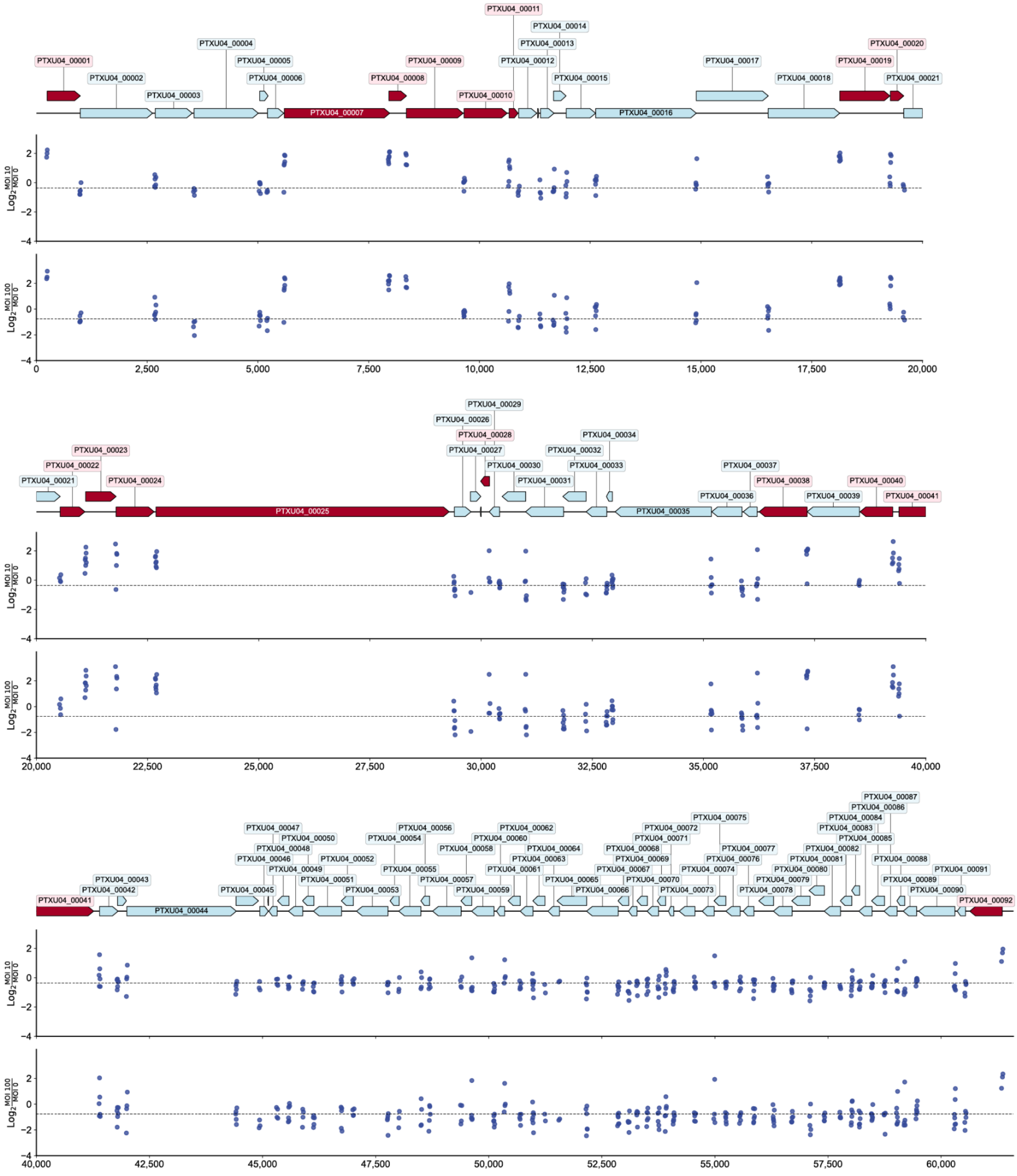


**Fig. S23. Genomewide ddCas13d-Fitness for Phage PTXU04**

(Top track) Gene overview of phage PTXU04 (MK373772). Coding genes are shown in light blue and highlighted in red if they met significance thresholds (Methods). tRNAs are shown in orange. Other noncoding genetic elements are shown in white. (Middle track) ddCas13d gene knockdown fitness when targeting PTXU04 genes during infection at 10 MOI. Median guide fitness is shown with a dashed line. (Bottom track) ddCas13d gene knockdown fitness when targeting PTXU04 genes during infection at 100 MOI. Median guide fitness is shown with a dashed line.

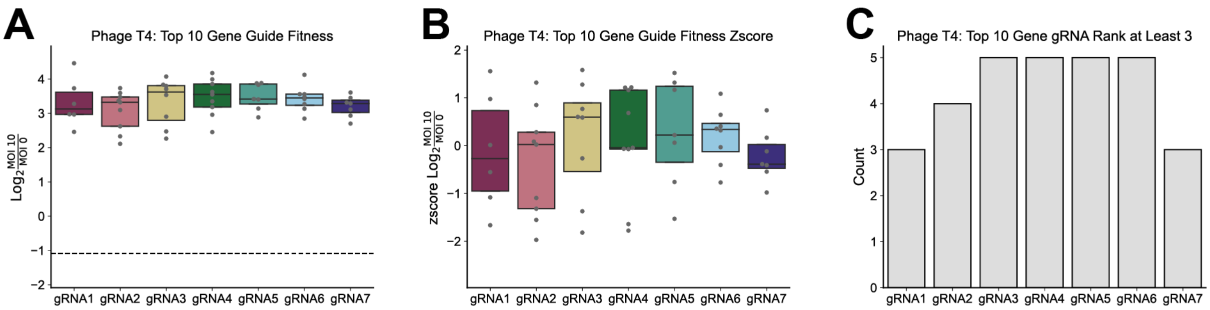


**Fig. S24. Guide Fitness Distributions for Top 10 Fit T4 Genes**

All analyses in fig. S24 refer to guides that target the Top 10 Fit Genes in T4 and pass quality control criteria (Methods). Genes with under 5 quality-control-passing guides were excluded. (**A**). Guide fitness distribution separated by relative gRNA position. Dashed line reflects median guide fitness observed in the experiments. (**B**). Z-score normalized guide fitness distribution (by targeted gene) separated by relative gRNA position. (**C**). Number of guides performing in the Top 3 guides for a given gene. Central line in boxplots in (**A**) and (**B**) represent the median and edges of the boxplot represents the 25 and 75 confidence intervals.

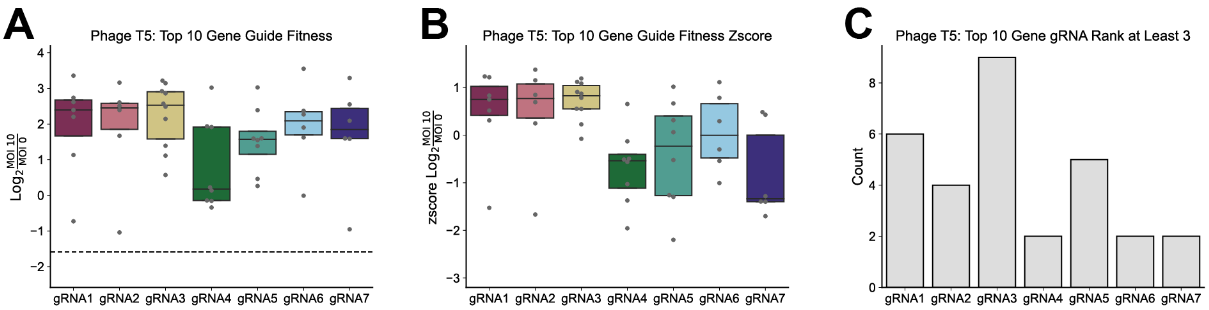


**Fig. S25. Guide Fitness Distributions for Top 10 Fit Phage T5 Genes**

All analyses in fig. S25 refer to guides that target the Top 10 Fit Genes in phage T5 and pass quality control criteria (Methods). Genes with under 5 quality-control-passing guides were excluded. (**A**). Guide fitness distribution separated by relative gRNA position. Dashed line reflects median guide fitness observed in the experiments. (**B**). Z-score normalized guide fitness distribution (by targeted gene) separated by relative gRNA position. (**C**). Number of guides performing in the Top 3 guides for a given gene. Central line in boxplots in (**A**) and (**B**) represent the median and edges of the boxplot represents the 25 and 75 confidence intervals.

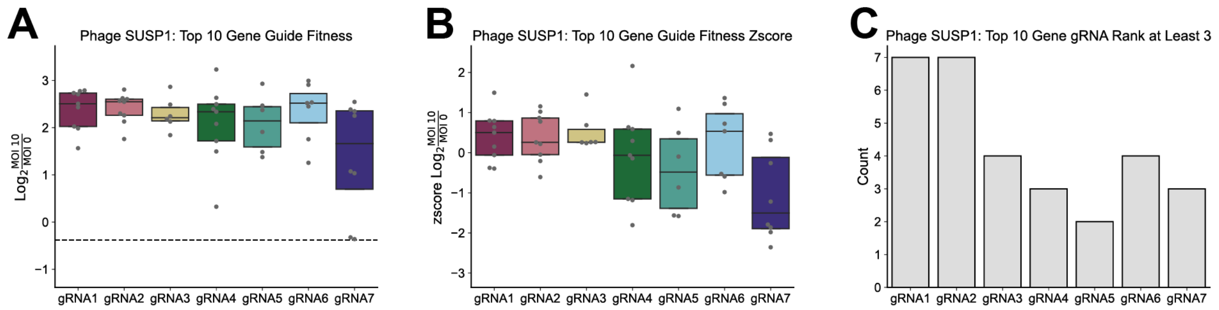


**Fig. S26. Guide Fitness Distributions for Top 10 Fit Phage SUSP1 Genes**

All analyses in fig. S26 refer to guides that target the Top 10 Fit Genes in phage SUSP1 and pass quality control criteria (Methods). Genes with under 5 quality-control-passing guides were excluded. (**A**). Guide fitness distribution separated by relative gRNA position. Dashed line reflects median guide fitness observed in the experiments. (**B**). Z-score normalized guide fitness distribution (by targeted gene) separated by relative gRNA position. (**C**). Number of guides performing in the Top 3 guides for a given gene. Central line in boxplots in (**A**) and (**B**) represent the median and edges of the boxplot represents the 25 and 75 confidence intervals.

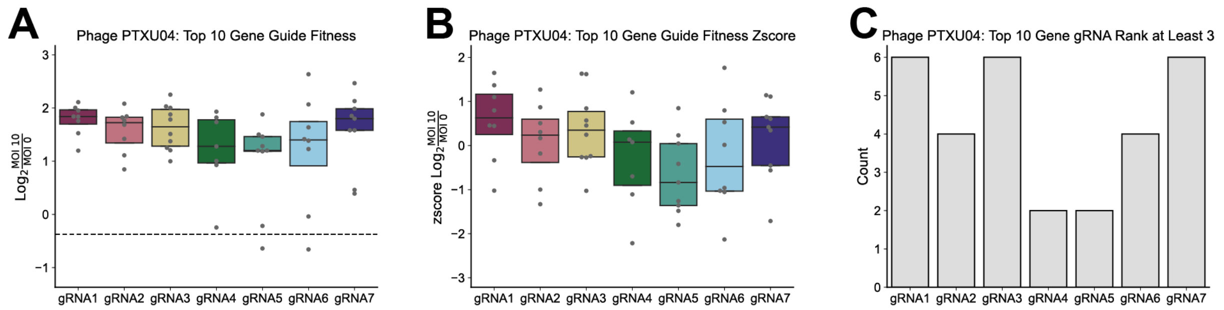


**Fig. S27. Guide Fitness Distributions for Top 10 Fit Phage PTXU04 Genes**

All analyses in fig. S27 refer to guides that target the Top 10 Fit Genes in phage PTXU04 and pass quality control criteria (Methods). Genes with under 5 quality-control-passing guides were excluded. (**A**). Guide fitness distribution separated by relative gRNA position. Dashed line reflects median guide fitness observed in the experiments. (**B**). Z-score normalized guide fitness distribution (by targeted gene) separated by relative gRNA position. (**C**). Number of guides performing in the Top 3 guides for a given gene. Central line in boxplots in (**A**) and (**B**) represent the median and edges of the boxplot represents the 25 and 75 confidence intervals.

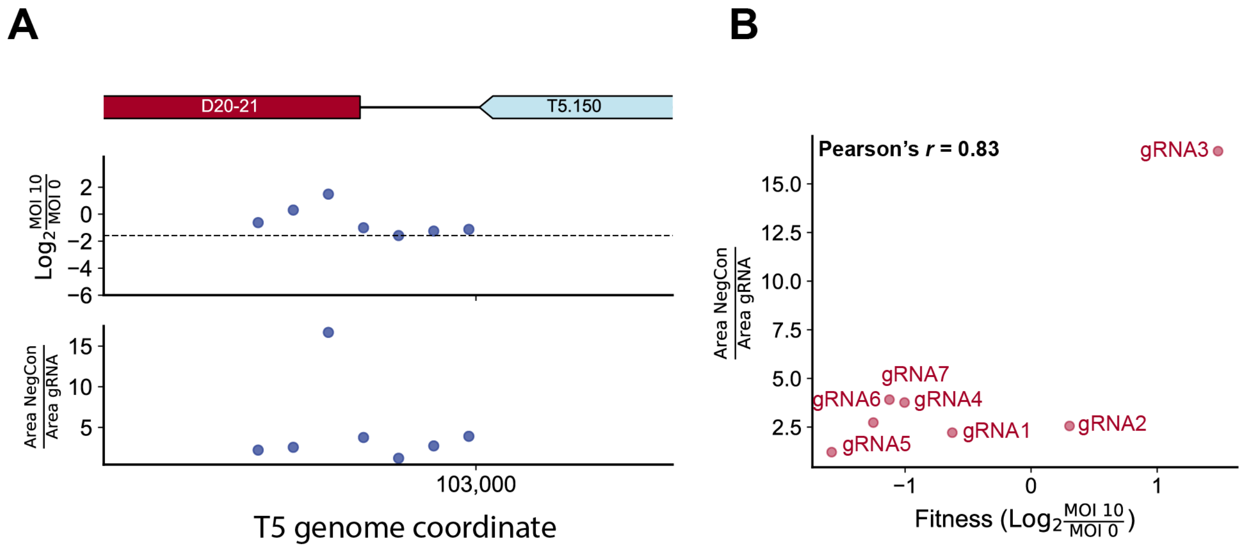


**Fig. S28. Guide fitness versus plaque size for T5 *D20-21* (*mcp*)**

(**A**). Fitness and fold plaque size reduction for CRISPRi-ART targeting of T5 *D20-21* (*mcp*) by position. Plots show gene organization at the *D20-21* locus (top), fitness values from pooled library screens (middle), and fold plaque size reduction from individually tested crRNAs (bottom). (**B**). Correlation between fitness values from pooled library screens and fold plaque size reduction. Correlation is reported with Pearson's coefficient (*r*).
